# The mycotoxin zearalenone couples secondary metabolism to *N*-acetylglucosamine catabolism

**DOI:** 10.64898/2026.11.04.764435

**Authors:** Nahyun Lee, Juwon Yang, Jiyeun Park, Jae Woo Han, Hun Kim, Hokyoung Son

## Abstract

Fungal secondary metabolites are often viewed as terminal products of specialized metabolism, yet how their production influences the physiology of the producing organism remains poorly understood. Here, we show that zearalenone (ZEA) production is functionally coupled to *N*-acetylglucosamine (GlcNAc) catabolism in *Fusarium graminearum*. Transcriptome profiling of a *ZEB2*-overexpressing strain revealed enrichment of GlcNAc metabolic functions, and exogenous ZEA induced key catabolic genes independently of Zeb2. Among pathway components, *NAG*, encoding GlcNAc-6-phosphate deacetylase, was critical for GlcNAc utilization, and its loss caused severe GlcNAc-dependent growth inhibition. Genetic impairment of *NAG* was poorly tolerated under elevated ZEA pathway activity. Conversely, *NAG* overexpression increased tolerance to exogenous ZEA and alleviated the growth defect associated with *ZEB2* overexpression. Together, these findings reveal a functional connection between secondary metabolite production and primary nutrient metabolism and suggest that ZEA-associated metabolic remodeling contributes to the fitness of the producing fungus.

## Introduction

Fungal secondary metabolites have been extensively studied for their diverse biological activities and ecological roles, including competition, defense, and interactions with other organisms ^1–4^. Although considerable progress has been made in elucidating their biosynthetic pathways and regulatory mechanisms, their physiological significance within the fungi that produce them remains a longstanding question in fungal biology. Increasing evidence, however, indicates that some specialized metabolites also influence the physiology of the producing organism by modulating development, stress adaptation, or metabolic homeostasis ^2,3,5,6^. For example, gliotoxin production in *Aspergillus fumigatus* is tightly coupled to self-protection mechanisms involving sulfur metabolism and oxidative stress defense, thereby limiting the detrimental effects associated with endogenous toxin accumulation ^7^. Other fungal metabolites participate in broader regulatory processes: butyrolactone I in *A. terreus* coordinates secondary metabolism with cellular metabolic activity to promote lovastatin production ^8^, whereas the quorum-sensing molecule farnesol in *Candida albicans* regulates morphogenesis, oxidative stress responses, and drug resistance ^9–11^. Aflatoxin production in *A. flavus* has also been associated with oxidative stress adaptation under specific conditions ^12^. Despite these advances, how fungal secondary metabolites influence primary metabolic processes and contribute to the physiology of their producers remains poorly understood.

Among mycotoxin-producing phytopathogens, *Fusarium graminearum* is a major causal agent of Fusarium head blight in small-grain cereals and Gibberella ear rot in maize, resulting in substantial yield losses and reduced grain quality ^13^. Several *Fusarium* species, including *F. graminearum*, *F. culmorum*, *F. equiseti*, and *F. pseudograminearum*, produce zearalenone (ZEA), a well-characterized estrogenic polyketide mycotoxin that contaminates agricultural commodities and poses risks to animal and human health ^14,15^. ZEA biosynthesis is controlled by a dedicated gene cluster containing *PKS4*, *PKS13*, *ZEB1*, and the bZIP transcription factor *ZEB2* ^16,17^. ZEA production is further regulated by two Zeb2 isoforms, the transcriptionally active Zeb2L and the inhibitory Zeb2S, which together form a tightly controlled feedback system ^18^. Despite extensive knowledge of ZEA biosynthesis and its toxicological effects, considerably less is known about how ZEA production influences the physiology and metabolic state of the producing fungus.

Our previous transcriptome-based analysis of Zeb2-responsive genes revealed that activation of the ZEA regulatory network extends beyond the canonical biosynthetic cluster and includes genes involved in diverse cellular processes. This observation prompted us to examine whether elevated ZEA pathway activity is accompanied by coordinated changes in primary metabolism. *N*-acetylglucosamine (GlcNAc) is the fundamental building block of fungal chitin and can serve as an alternative source of carbon and nitrogen ^19^. In dimorphic fungi such as *C. albicans*, GlcNAc catabolism is closely associated with morphogenetic signaling ^20–22^, whereas in filamentous fungi it contributes to nutrient utilization, development, and adaptation to nutrient availability ^23–27^. However, whether and how GlcNAc metabolism is coordinated with mycotoxin production remains poorly understood. Here, we investigated the relationship between ZEA production and GlcNAc catabolism. Our transcriptomic and genetic analyses revealed that ZEA induces GlcNAc catabolic gene expression and that *NAG*, encoding GlcNAc-6-phosphate deacetylase, contributes to fungal fitness under elevated ZEA pathway activity. Together, these findings uncover a previously unrecognized coupling between fungal secondary metabolism and amino-sugar utilization and suggest that ZEA-associated remodeling of primary nutrient metabolism contributes to physiological adaptation in the producing fungus.

## Results

### Transcriptome analysis reveals enrichment of GlcNAc metabolism in the *ZEB2*-overexpressing strain

Our previous comparative transcriptome analysis of the wild-type and *ZEB2*-overexpressing strains identified 393 genes that were upregulated in *ZEB2OE* (log₂ fold change ≥ 2) (Supplementary Data 1). To characterize the biological processes associated with *ZEB2* overexpression, these genes were subjected to Gene Ontology (GO) enrichment and Kyoto Encyclopedia of Genes and Genomes (KEGG) pathway analyses (Fig. 1).

**Fig. 1.**
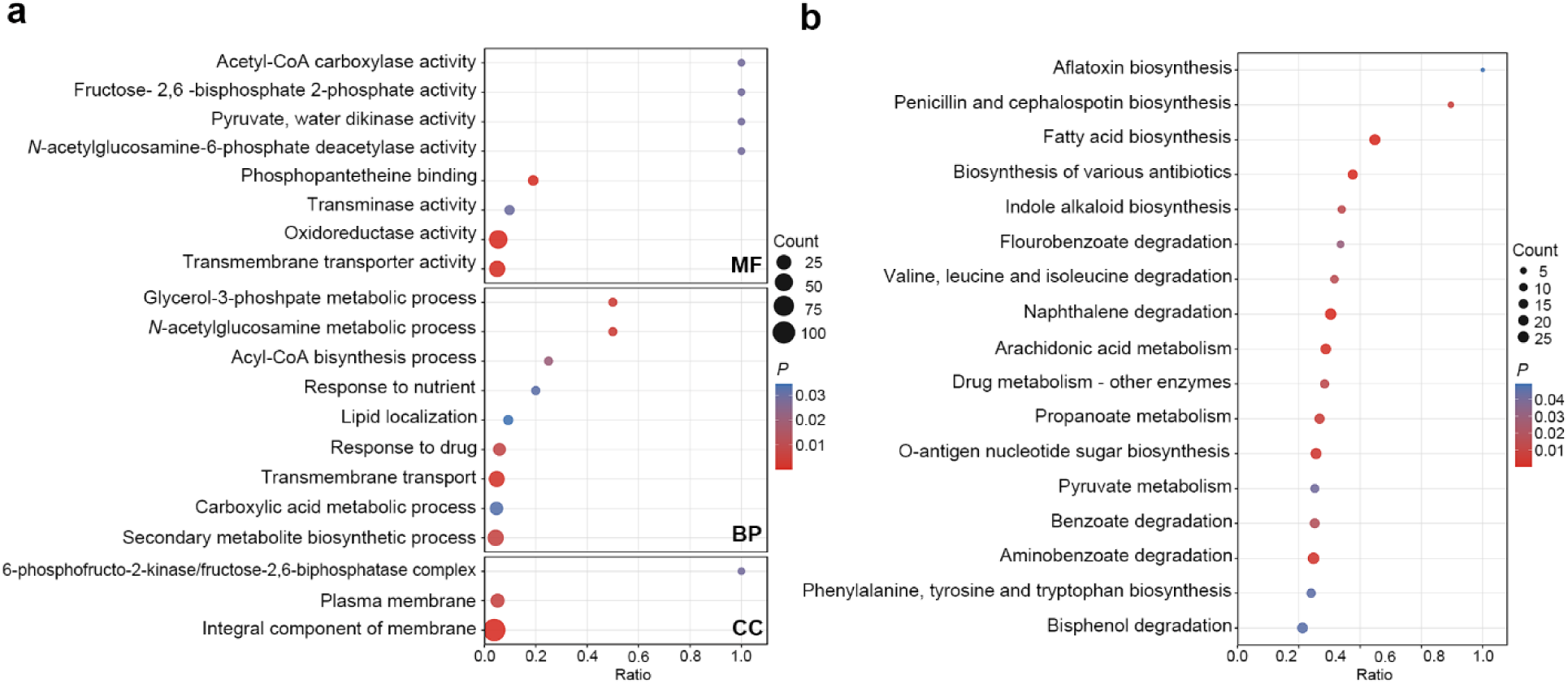
Functional enrichment analysis of genes upregulated in the *ZEB2OE* strain. **a** Gene Ontology (GO) enrichment analysis of 393 genes upregulated in *ZEB2OE* relative to the wild type (log_2_ fold change ≥ 2). Enriched GO terms are classified into molecular function (MF), biological process (BP), and cellular component (CC) categories. **b** Kyoto Encyclopedia of Genes and Genomes (KEGG) pathway enrichment analysis of the same gene set. In both panels, dot size represents the number of genes associated with each term or pathway (Count), dot color indicates the *P* value, and the x-axis represents the enrichment ratio. Only significantly enriched terms and pathways (*P* < 0.05) are shown.

GO enrichment analysis identified enrichment of multiple metabolic and transport-related functions (Fig. 1a). Within the Biological Process (BP) category, enriched terms included secondary metabolite biosynthetic process, transmembrane transport, response to drug, carboxylic acid metabolic process, and *N*-acetylglucosamine (GlcNAc) metabolic process. The Molecular Function (MF) category was enriched for oxidoreductase activity and transmembrane transporter activity, together with several metabolic enzyme activities, including *N*-acetylglucosamine-6-phosphate deacetylase activity. In the Cellular Component (CC) category, terms associated with the plasma membrane and integral component of membrane were prominently represented.

KEGG pathway analysis further revealed enrichment of multiple metabolic pathways in the *ZEB2OE* strain, including pathways associated with secondary metabolism, lipid metabolism, and central carbon metabolism (Fig. 1b). These pathways encompassed various antibiotic biosynthetic pathways, fatty acid biosynthesis, drug metabolism, propanoate metabolism, and pyruvate metabolism. Enrichment of O-antigen nucleotide sugar biosynthesis was also observed, suggesting alterations in sugar-related metabolic processes.

Notably, the GlcNAc metabolic process was significantly enriched despite being represented by a relatively small number of genes (Fig. 1a). Moreover, *N*-acetylglucosamine-6-phosphate deacetylase activity, a key enzymatic function in GlcNAc catabolism, was independently identified among the enriched MF terms. Whereas the enrichment of secondary metabolite-related functions was consistent with the established role of Zeb2 in ZEA biosynthesis, the concurrent enrichment of GlcNAc-related functions suggested an unexpected connection between ZEA pathway activation and primary nutrient metabolism. We therefore focused subsequent analyses on the GlcNAc catabolic pathway to investigate its relationship with Zeb2-mediated ZEA production.

### ZEA induces GlcNAc catabolic gene expression independently of *ZEB2*

To identify the GlcNAc catabolic machinery in *F. graminearum*, we performed BLASTp searches using previously characterized GlcNAc catabolic proteins from *Neurospora crassa* as queries ^26^. Previous studies have shown that GlcNAc catabolism is mediated by a conserved set of proteins comprising the GlcNAc transporter Ngt, GlcNAc kinase Hxk, GlcNAc-6-phosphate deacetylase Nag, glucosamine-6-phosphate deaminase Dam, and the transcriptional regulator Ron. We identified putative homologs of these proteins based on sequence similarity and genomic organization (Supplementary Data 2). Notably, the corresponding *NGT*, *HXK*, and *NAG* candidates were also upregulated in the *ZEB2OE* transcriptome. Although the selected *DAM* candidate did not show the highest sequence identity, its strong upregulation in *ZEB2OE* and genomic proximity to *NAG*, *RON*, and *HXK* supported its identification as a putative homolog. Examination of the genomic organization of the selected genes revealed that *NAG*, *RON*, *DAM*, and *HXK* form a contiguous gene cluster, whereas the putative GlcNAc transporter gene *NGT* is located at a separate locus. This organization resembles the conserved GlcNAc catabolic gene cluster reported in other filamentous fungi ^26,28^. Consistent with their predicted roles, all five genes were induced by GlcNAc supplementation, with particularly strong responses observed for *NAG* and *DAM* (Supplementary Fig. 1).

To examine whether the expression of GlcNAc catabolic genes was associated with Zeb2 activity or ZEA production, we compared their transcript levels in the wild-type, Δ*zeb2*, and *ZEB2OE* strains. Under ZEA-inducing conditions, expression levels of *NGT*, *HXK*, *NAG*, and *DAM* were significantly higher in *ZEB2OE* than in the wild-type strain (Fig. 2). These results validated the transcriptomic findings and established an association between *ZEB2* overexpression and increased expression of GlcNAc catabolic genes.

**Fig. 2.**
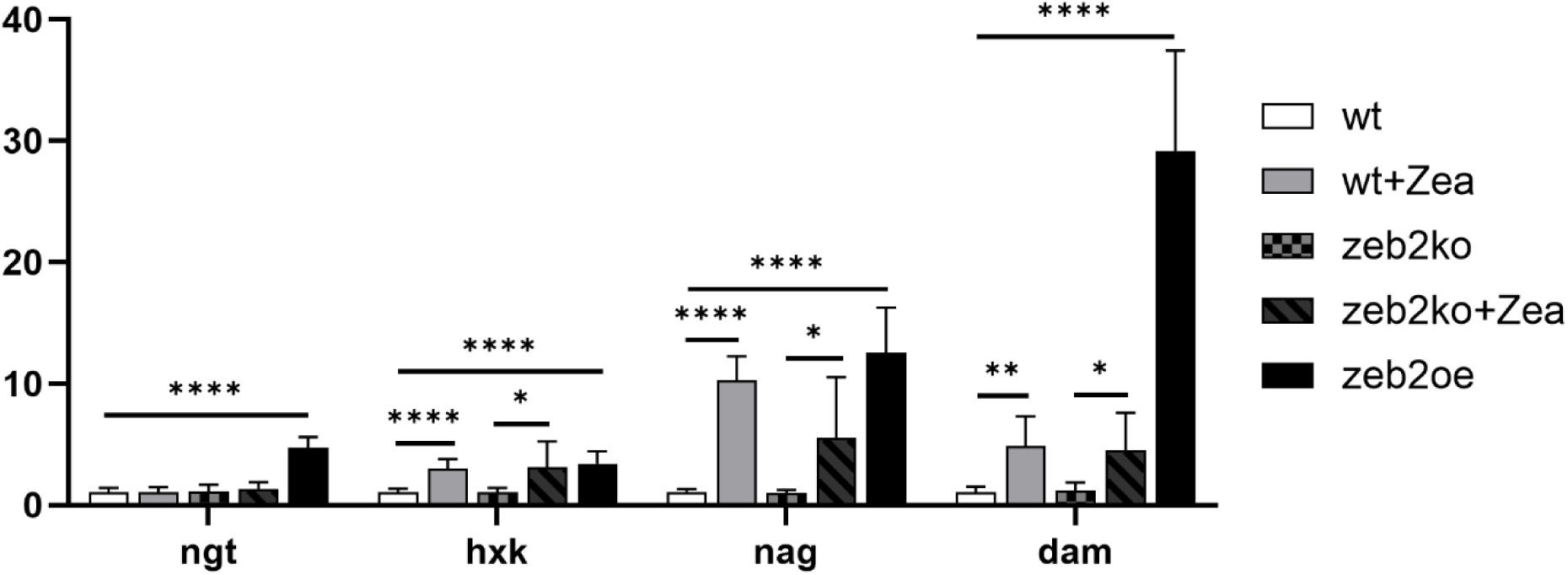
ZEA induces expression of key GlcNAc catabolic genes independently of Zeb2 in *F. graminearum*. Relative transcript levels of *RON*, *NGT*, *HXK*, *NAG*, and *DAM* were determined by RT-qPCR in the wild-type (WT), Δ*zeb2*, and *ZEB2OE* strains cultured in SG medium for 3 days. The wild-type and Δ*zeb2* strains were treated with 30 µM exogenous ZEA, as indicated. Transcript levels were normalized to *CYP1* and are presented relative to those in the untreated wild type. Data are presented as mean ± SD from three independent biological replicates. Statistical significance for the indicated comparisons was determined using Welch’s *t*-test. ns, not significant; \**P* < 0.05; \*\**P* < 0.01; and \*\*\*\**P* < 0.0001.

We next asked whether ZEA itself contributed to the induction of these genes. Treatment of the wild-type strain with 30 µM exogenous ZEA markedly increased the expression of *HXK*, *NAG*, and *DAM* compared with that in the untreated wild-type control, with particularly pronounced increases in *NAG* and *DAM* expression (Fig. 2). However, because ZEA also induces the transcriptionally active Zeb2L isoform through a previously characterized feedback mechanism ^18^, it remained unclear whether the observed induction of GlcNAc catabolic genes required Zeb2.

To distinguish between these possibilities, we examined the response of the Δ*zeb2* mutant to exogenous ZEA. Despite the absence of *ZEB2*, ZEA treatment significantly increased the expression of key GlcNAc catabolic genes compared with the untreated Δ*zeb2* mutant (Fig. 2). These results demonstrate that ZEA induces a GlcNAc catabolic transcriptional response independently of Zeb2, revealing an unexpected regulatory connection between a fungal mycotoxin and primary nutrient metabolism. In contrast, *RON* did not exhibit a significant or consistent increase under the tested conditions, suggesting that its transcriptional regulation may differ from that of the other GlcNAc catabolic genes.

To assess whether the GlcNAc response was broadly associated with mycotoxin exposure, we next examined whether deoxynivalenol (DON), another major *Fusarium* mycotoxin, elicited a similar transcriptional response. The wild-type strain was cultured under DON-inducing conditions with or without 3 µM exogenous DON, and *NAG* transcript levels were measured at 3 and 7 days. Exogenous DON did not significantly alter *NAG* expression at either time point, with transcript levels remaining comparable between DON-treated and untreated cultures (Supplementary Fig. 2). Unlike ZEA, DON did not induce *NAG* expression under the conditions tested, suggesting that the ZEA-associated transcriptional response is not a universal consequence of mycotoxin exposure.

### Loss of *NAG* causes severe GlcNAc-dependent growth inhibition

To investigate the physiological roles of the GlcNAc catabolic genes, deletion mutants were evaluated for radial growth on complete medium (CM), carbon-free minimal medium (MM−C), and MM−C supplemented with GlcNAc, glucose, or both (Fig. 3a, b). All strains showed comparable growth on CM, whereas distinct growth phenotypes emerged under minimal and carbon-supplemented conditions. Among the GlcNAc catabolic mutants, Δ*nag* exhibited the most pronounced phenotype, displaying reduced radial growth on MM−C and almost complete growth inhibition on MM−C supplemented with GlcNAc. In comparison, Δ*ron* and Δ*hxk* showed moderate reductions in growth in the presence of GlcNAc. The Δ*ngt* mutant showed no apparent growth defect and instead exhibited slightly enhanced growth under GlcNAc-supplemented conditions, whereas Δ*dam* was largely unaffected by GlcNAc supplementation.

**Fig. 3.**
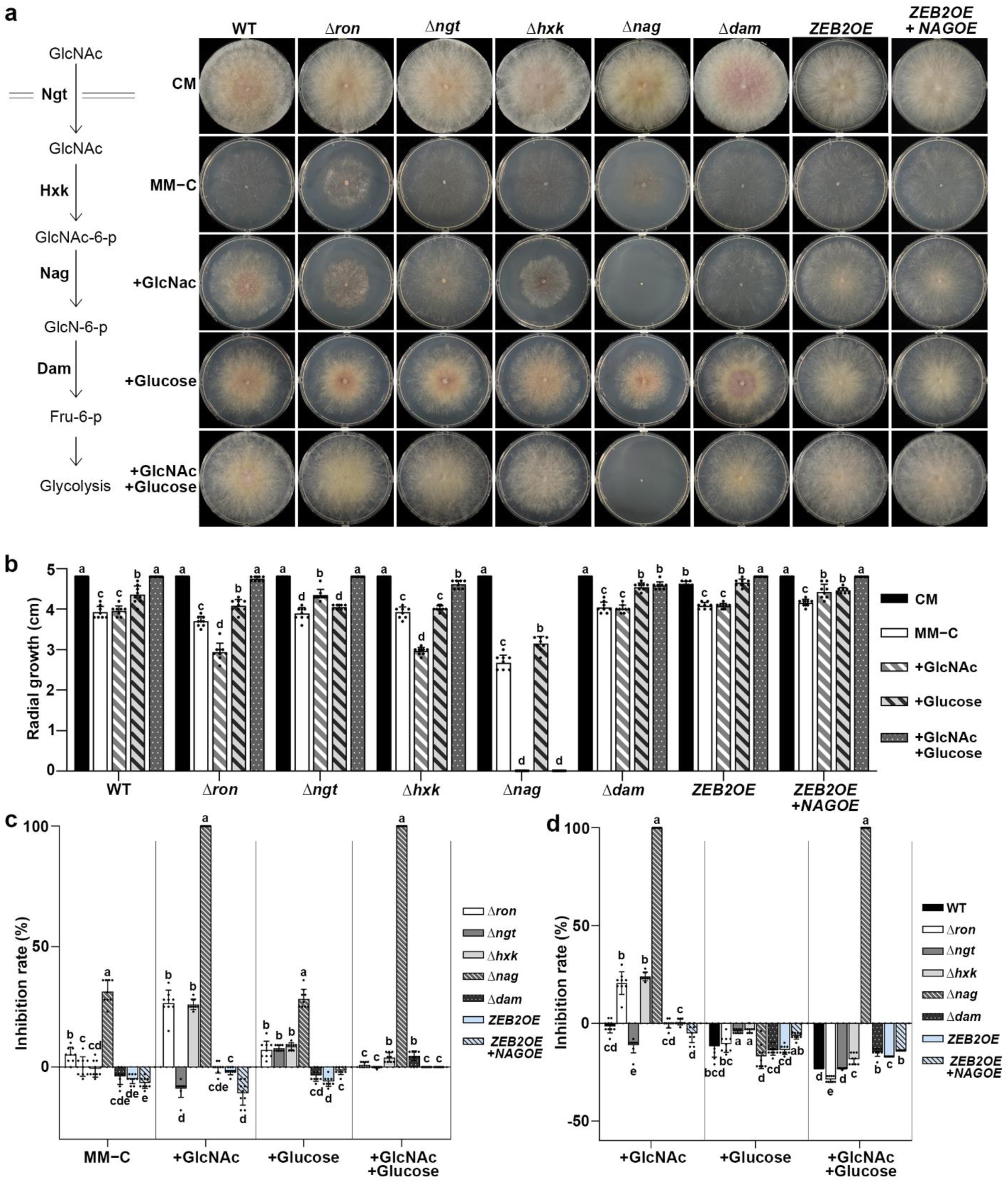
Loss of *NAG* causes severe GlcNAc-dependent growth inhibition in *F. graminearum*. **a** Schematic representation of the GlcNAc catabolic pathway and representative colony morphology of the indicated strains grown on complete medium (CM), carbon source-free minimal medium (MM−C), or MM−C supplemented with 1% (w/v) GlcNAc, 1% (w/v) glucose, or both for 3 days. **b** Radial growth of the indicated strains under the conditions shown in **a**. Different lowercase letters indicate significant differences among culture conditions within each strain, as determined by one-way ANOVA followed by Tukey’s HSD test (*P* < 0.05). **c** Growth inhibition of each strain relative to the wild type under the corresponding nutritional condition. Growth inhibition was calculated as [(radial growth of WT − radial growth of the indicated strain)/radial growth of WT] × 100. Negative values indicate greater growth than that of the wild type under the corresponding condition. **d** Growth inhibition under carbon-supplemented conditions relative to its growth on MM−C within each strain. The ratio was calculated as [(radial growth on MM−C − radial growth under the indicated supplemented condition)/radial growth on MM−C] × 100. Negative values indicate enhanced growth relative to MM−C. For **c** and **d**, different lowercase letters indicate significant differences among strains within each nutritional condition, as determined by one-way ANOVA followed by Tukey’s HSD test (*P* < 0.05). Data are presented as mean ± SD from independent biological replicates.

To compare mutant growth directly with that of the wild type under each nutritional condition, growth inhibition was calculated relative to the wild type (Fig. 3c). The Δ*nag* mutant showed approximately 32% growth inhibition on MM−C, which increased to nearly 100% upon GlcNAc supplementation. Although supplementation of glucose supported growth of Δ*nag*, the simultaneous addition of GlcNAc and glucose again resulted in almost complete growth inhibition. Thus, the severe GlcNAc-dependent phenotype of Δ*nag* was not alleviated by the availability of an alternative utilizable carbon source. To distinguish the effects of carbon supplementation from differences in basal growth among mutants, we also normalized growth to each strain’s performance on MM−C (Fig. 3d). This analysis confirmed that GlcNAc exerted a particularly strong inhibitory effect on Δ*nag*, whereas glucose supported its growth.

To further determine whether the Δ*nag* phenotype was specifically dependent on GlcNAc, the wild-type and Δ*nag* strains were grown on glucose-containing MM−C supplemented with increasing concentrations of GlcNAc. Growth inhibition of Δ*nag* became apparent at 0.01% GlcNAc and became progressively more severe at higher concentrations (Supplementary Fig. 3). Importantly, the GlcNAc-dependent growth defect was rescued by *NAG* complementation or overexpression, confirming that the phenotype resulted specifically from loss of *NAG* function. As an additional control, we examined a Δ*chs7* mutant lacking chitin synthase Chs7 and observed enhanced growth following GlcNAc supplementation ^29^. This contrasting response further supports the conclusion that the pronounced GlcNAc sensitivity of Δ*nag* results from impaired *NAG*-dependent metabolism rather than a general inhibitory effect of GlcNAc. Consistent with the radial growth phenotype, a PrestoBlue-based assay showed a pronounced concentration-dependent decrease in cellular metabolic activity in Δ*nag*, whereas the wild type maintained substantially greater metabolic activity across the tested GlcNAc concentrations (Supplementary Fig. 4). Together, these results demonstrate that the severe phenotype of Δ*nag* cannot be explained solely by an inability to utilize GlcNAc as a carbon source. Instead, disruption of the *NAG*-dependent step renders GlcNAc exposure strongly inhibitory, revealing a critical role for *NAG* in maintaining fungal fitness during amino-sugar utilization.

### *ZEB2* overexpression exacerbates growth defects caused by impaired *NAG* function

Based on the transcriptional association between *ZEB2* overexpression and GlcNAc metabolism, we hypothesized that enhanced *ZEB2* activity increases the cellular requirement for *NAG*-dependent GlcNAc metabolism. To test this hypothesis, we first attempted to generate a Δ*nag* mutant in the *ZEB2OE* background. However, no transformants carrying the desired genotype were recovered despite three independent transformation attempts. Similarly, outcrossing the *ZEB2OE* strain with the Δ*nag* mutant failed to yield progeny carrying both genetic modifications despite repeated crosses and subsequent progeny screening ^30^. These repeated failures are consistent with a strong negative genetic interaction between *NAG* deletion and *ZEB2* overexpression.

To assess the consequences of reduced NAG function in a genetically tractable background, we generated a conditional expression strain in which *NAG* was placed under the control of the copper-responsive *FCR1* promoter (*P_FCR1_-FgNAG*) ^31^. The conditional strain was evaluated on MM−C supplemented with either 1% glucose or 1% GlcNAc in the presence of CuSO₄ or the copper chelator bathocuproinedisulfonic acid (BCS) (Fig. 4a). On glucose-containing medium, the *P_FCR1_-FgNAG* strain maintained substantial growth under the tested conditions (Fig. 4b). In contrast, on GlcNAc-containing medium, the conditional strain exhibited pronounced growth inhibition under both basal and CuSO₄-treated conditions, with inhibition rates of approximately 70–75% relative to the wild type (Fig. 4b). BCS supplementation largely restored growth, consistent with relief of copper-mediated repression of *NAG*.

**Fig. 4.**
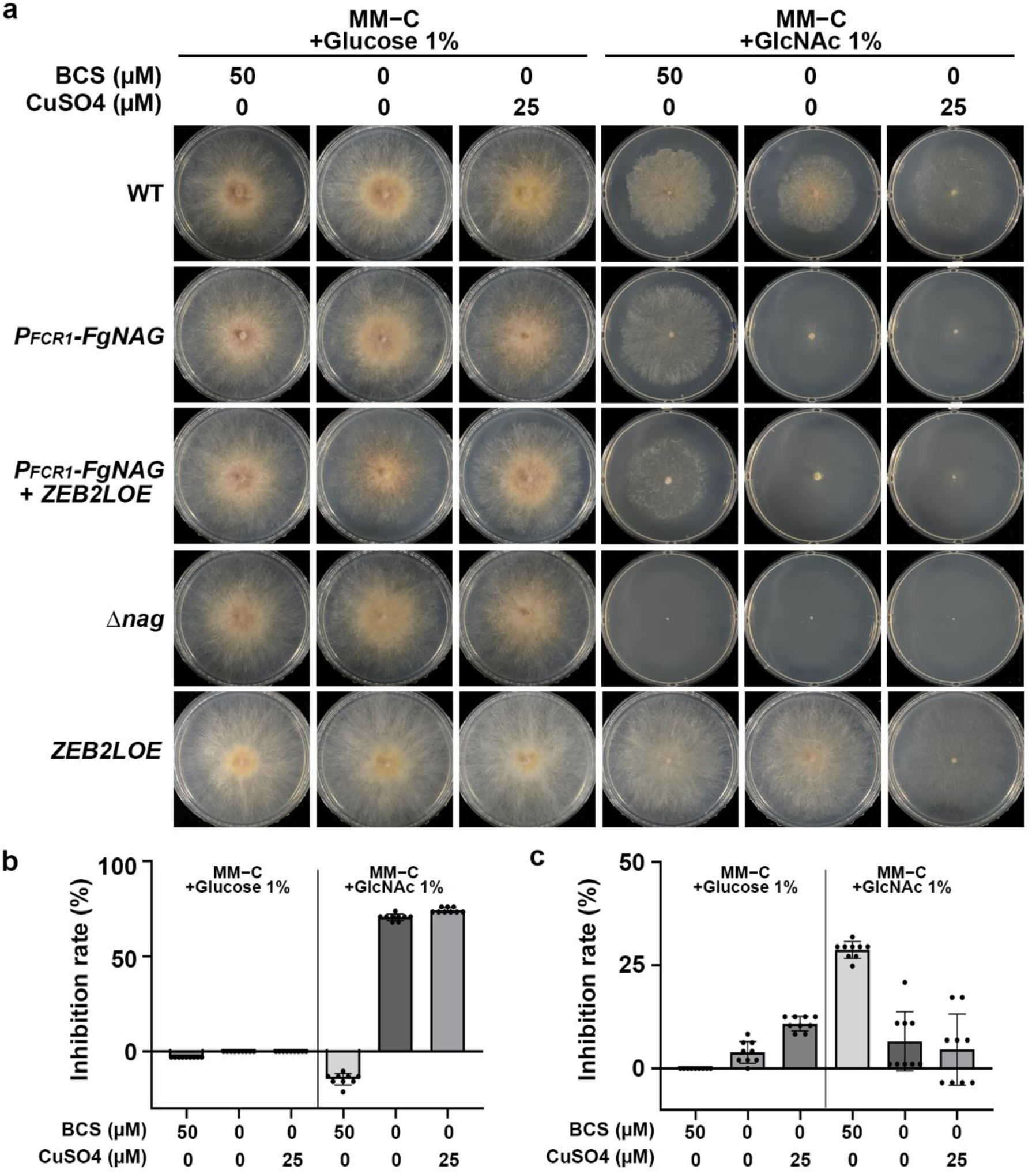
Conditional modulation of *FgNAG* reveals a genetic interaction with *ZEB2L* overexpression. **a** Representative colony morphology of the wild-type, *P_FCR1_-FgNAG*, *P_FCR1_-FgNAG*; *ZEB2LOE*, Δ*nag*, and *ZEB2LOE* strains grown on carbon source-free minimal medium (MM−C) supplemented with 1% (w/v) glucose or 1% (w/v) GlcNAc. Conditional expression of *FgNAG* from the copper-responsive *FCR1* promoter was modulated by supplementation with 50 µM BCS or 25 µM CuSO₄. CuSO₄ represses *FgNAG* expression, whereas BCS relieves copper-dependent repression. **b** Growth inhibition of *P_FCR1_-FgNAG* relative to the wild type under the corresponding culture conditions. **c** Growth inhibition of *P_FCR1_-FgNAG*; *ZEB2LOE* relative to the parental *P_FCR1_-FgNAG* strain under each culture condition. Growth inhibition was calculated as [(radial growth of the reference strain − radial growth of the indicated strain)/radial growth of the reference strain] × 100. Negative values indicate greater growth than that of the corresponding reference strain. Data are presented as mean ± SD from three independent biological replicates.

We then introduced a *ZEB2L* overexpression construct into the *P_FCR1_-FgNAG* background, and elevated *ZEB2* expression was confirmed by reverse transcription quantitative PCR (RT-qPCR) (Supplementary Fig. 5). On glucose-containing medium, the *P_FCR1_-FgNAG*; *ZEB2LOE* strain exhibited greater growth inhibition than the parental *P_FCR1_-FgNAG* strain, whereas BCS supplementation largely restored growth (Fig. 4a, c). On GlcNAc-containing medium, both conditional strains showed severe growth inhibition under basal and CuSO_4_-treated conditions, precluding a reliable assessment of the additional effect of *ZEB2L* overexpression. In contrast, under BCS treatment, the *P_FCR1_-FgNAG*; *ZEB2LOE* strain remained approximately 25–30% inhibited relative to the parental strain (Fig. 4c). Together with the repeated inability to recover a Δ*nag*; *ZEB2OE* double mutant, these results reveal a strong genetic interaction between *ZEB2* overexpression and *NAG* function, highlighting the critical importance of *NAG*-dependent metabolism under elevated ZEA pathway activity.

### *NAG* overexpression alleviates ZEA-associated cellular stress

Given the strong genetic interaction between *ZEB2* overexpression and *NAG*, we investigated whether increasing *NAG* expression could protect the fungus against ZEA exposure ^32^. The wild-type and two independent *NAG*-overexpressing strains (*NAGOE*) were exposed to increasing concentrations of exogenous ZEA (0–80 µM), and cellular metabolic activity was measured at 2 days post-inoculation using the PrestoBlue assay (Fig. 5a, b). ZEA caused a concentration-dependent increase in inhibition in the wild-type strain, reaching nearly complete inhibition at higher concentrations. In contrast, both *NAGOE* strains exhibited lower inhibition rates than the wild type, with the protective effect being particularly evident at intermediate to high ZEA concentrations. These results indicate that *NAG* overexpression increases fungal tolerance to exogenous ZEA.

**Fig. 5.**
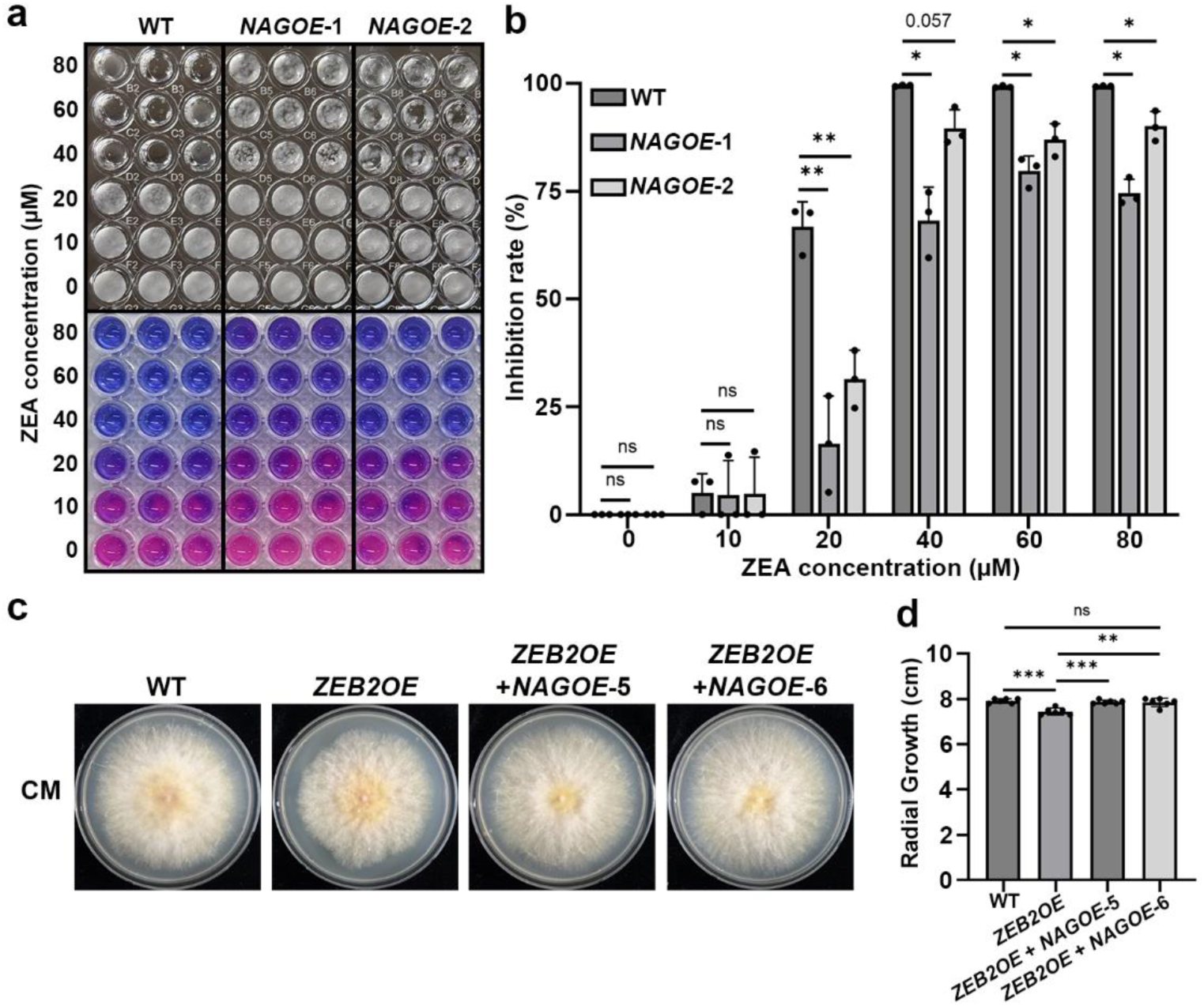
*NAG* overexpression enhances tolerance to ZEA and alleviates the growth defect associated with *ZEB2* overexpression. **a** Representative cultures and PrestoBlue reactions of the wild-type and two independent *NAG*-overexpressing strains exposed to increasing concentrations of exogenous ZEA (0–80 µM). Cellular metabolic activity was assessed at 2 days post-inoculation using the PrestoBlue assay. **b** Inhibition rates of the strains shown in **a** relative to the corresponding untreated controls. **c** Representative colony morphology of the wild-type, *ZEB2OE*, and two independent *ZEB2OE*; *NAGOE* strains grown on complete medium (CM). **d** Quantification of radial growth of the strains shown in **c**. Data are presented as mean ± SD from three independent biological replicates. Statistical significance was determined using Welch’s two-tailed *t*-test for the indicated comparisons. \**P* < 0.05; \*\**P* < 0.01; ns, not significant.

We next examined whether increased *NAG* expression could alleviate the growth defect associated with constitutive ZEA pathway activation. Introduction of the *NAG* overexpression construct into the *ZEB2OE* background substantially restored radial growth, with independent *ZEB2OE*; *NAGOE* strains approaching wild-type growth levels (Fig. 5c, d). *NAG* overexpression also partially alleviated the GlcNAc-associated growth inhibition observed in the *ZEB2OE* strain (Fig. 3c, d). Together, these findings demonstrate that *NAG* overexpression mitigates the physiological burden associated with elevated ZEA pathway activity, supporting a protective role for GlcNAc catabolism in the producing fungus.

## Discussion

Despite decades of intensive research into fungal secondary metabolism, the physiological significance of many secondary metabolites within their producing organisms remains incompletely understood ^2,3^. Here, we uncover an unexpected functional connection between ZEA production and GlcNAc catabolism in *F. graminearum*. Our findings show that ZEA is sufficient to induce key GlcNAc catabolic genes independently of Zeb2, while genetic analyses reveal a strong interaction between elevated Zeb2 activity and *NAG* function. Moreover, *NAG* overexpression enhanced tolerance to exogenous ZEA and substantially alleviated the growth defect associated with constitutive ZEA pathway activation. Together, these findings extend the physiological significance of ZEA beyond its established biological activities toward other organisms and identify GlcNAc catabolism as a potential metabolic adaptation associated with the ZEA-producing state.

A key finding of this study is that ZEA itself induces GlcNAc catabolic gene expression independently of Zeb2. Previous studies have established that ZEA enhances the expression of the transcriptionally active Zeb2L isoform through a positive-feedback mechanism that promotes its own biosynthesis ^18^. However, the persistence of GlcNAc catabolic gene induction in the Δ*zeb2* mutant demonstrates that this response cannot be explained solely by Zeb2-mediated transcriptional regulation. This finding raises the possibility that ZEA acts not merely as an end product of secondary metabolism but also as a metabolic cue that influences primary metabolic processes in the producing fungus. Interestingly, *RON*, a putative transcriptional regulator of GlcNAc catabolism, did not exhibit a comparable response to ZEA, suggesting that ZEA-responsive regulation may differ from the canonical GlcNAc-induced transcriptional program. Rather than representing a passive consequence of *ZEB2* overexpression, the induction of GlcNAc catabolic genes may therefore reflect a distinct metabolic response associated with ZEA production, revealing an unexpected regulatory connection between fungal secondary metabolism and primary nutrient utilization.

The genetic interaction between *ZEB2* and *NAG* provides evidence that the connection between ZEA production and GlcNAc metabolism is functional rather than merely transcriptional. The repeated inability to recover a Δ*nag*; *ZEB2OE* double mutant, together with the enhanced growth defect observed when *ZEB2L* overexpression was combined with conditional restriction of *NAG* function, supports a strong functional interaction between these two processes. Conversely, increasing *NAG* expression enhanced tolerance to exogenous ZEA and alleviated growth defects associated with *ZEB2* overexpression. These reciprocal genetic effects therefore indicate that sufficient *NAG* function becomes particularly important under elevated ZEA pathway activity. Importantly, these findings do not demonstrate that Zeb2 directly increases GlcNAc catabolic flux. Rather, elevated ZEA pathway activity appears to establish a metabolic state in which efficient *NAG*-dependent metabolism becomes increasingly important for maintaining cellular fitness.

The physiological basis for this ZEA-associated activation of GlcNAc catabolism is likely multifactorial. One possibility is that functional GlcNAc metabolism contributes to tolerance of the physiological burden associated with ZEA accumulation. In *Aspergillus fumigatus*, gliotoxin self-protection engages redox, sulfur, and other homeostatic pathways, illustrating how production of a bioactive secondary metabolite can require compensatory remodeling of cellular physiology ^33^. In support of a related adaptive role in *F. graminearum*, *NAG* overexpression increased tolerance to exogenous ZEA and partially alleviated the growth defects of the *ZEB2OE* strain.

A second, complementary possibility is that GlcNAc utilization provides a metabolic compensation mechanism during elevated secondary metabolite production. Specialized metabolite biosynthesis can impose substantial demands on central carbon and redox metabolism; for example, production of the polyketide 6-methylsalicylic acid in *Aspergillus nidulans* competes with biomass formation for available acetyl-CoA, whereas penicillin production in *Penicillium chrysogenum* places a major burden on cytosolic NADPH supply ^34,35^. Given that ZEA is likewise a polyketide-derived metabolite, recruitment of an alternative nutrient-utilization pathway may help buffer the energetic and biosynthetic demands associated with elevated ZEA production.

A third, complementary possibility is that GlcNAc catabolism contributes to nutrient recycling during later stages of fungal growth. In *A. nidulans*, carbon starvation induces autolysis accompanied by the release of glucose and GlcNAc ^36,37^, while studies in other filamentous fungi have shown that chitin and chitosan turnover under carbon-limited conditions contributes to recycling of fungal cell-wall material ^38,39^. Nutrient recycling through autophagy is also important in *F. graminearum*, where endogenous cellular resources are mobilized under nutrient limitation to support fungal growth, development, and secondary metabolism ^40^. Such recycling could be particularly advantageous during later stages of fungal growth, when readily available nutrients decline and ZEA production becomes prominent. These mechanisms are not mutually exclusive and may together contribute to maintaining cellular fitness during elevated ZEA production.

At the pathway level, the GlcNAc catabolic machinery of *F. graminearum* is broadly conserved with that of other filamentous fungi, with *NAG*, *RON*, *DAM*, and *HXK* organized as a contiguous genomic cluster and the putative transporter *NGT* located at a separate locus. Similar organization has been reported in *Trichoderma reesei*, *Neurospora crassa*, and other fungi ^23,25,28^, supporting the evolutionary conservation of GlcNAc utilization as a coordinated metabolic module. Despite this structural conservation, however, the physiological roles of individual pathway components differed substantially in *F. graminearum*. Among the mutants examined, loss of *NAG* produced the most pronounced GlcNAc-dependent phenotype, whereas disruption of *RON* or *HXK* caused only moderate defects and deletion of *NGT* or *DAM* had relatively minor effects. This contrasts with other fungal systems in which disruption of several GlcNAc catabolic components substantially compromises growth on GlcNAc, suggesting that the relative contribution of individual pathway components may vary among fungal species ^25,28,41,42^. Thus, although the pathway architecture is conserved, *NAG* appears to represent a particularly critical metabolic node for GlcNAc utilization in *F. graminearum*.

The severe phenotype of Δ*nag* cannot be explained simply by an inability to use GlcNAc as a carbon source. Glucose supported growth of Δ*nag*, whereas GlcNAc strongly inhibited growth even in the presence of glucose as an alternative utilizable carbon source. This phenotype became progressively more severe with increasing GlcNAc concentrations and was restored by *NAG* complementation. Moreover, GlcNAc supplementation enhanced rather than inhibited growth of Δ*chs7*, arguing against intrinsic GlcNAc toxicity. A plausible explanation is that loss of Nag-mediated deacetylation leads to the accumulation of GlcNAc-6-phosphate upstream of the blocked metabolic step. A similar phenotype has been reported in *Candida albicans*, where deletion of the GlcNAc-6-phosphate deacetylase *DAC1* caused severe GlcNAc-dependent growth inhibition ^43^. Likewise, disruption of GlcNAc uptake or phosphorylation alleviated GlcNAc-induced growth inhibition in *Neurospora crassa* ^26^. Together, these findings suggest that the Δ*nag* phenotype may result from GlcNAc-6-phosphate accumulation and the associated metabolic imbalance.

The lack of additional *NAG* induction following exogenous DON treatment, in contrast to the pronounced response to ZEA, further suggests that the GlcNAc response is not simply a generic consequence of mycotoxin exposure. Nevertheless, comparison with a single additional mycotoxin is insufficient to establish strict ZEA specificity, and responses to other secondary metabolites will need to be examined. It also remains unclear whether ZEA is sensed directly or whether GlcNAc catabolic gene induction occurs in response to a downstream physiological consequence of ZEA accumulation. The molecular sensor or transcriptional regulator that links ZEA accumulation to GlcNAc catabolic gene expression is unknown, and direct measurements of GlcNAc metabolic flux and intracellular GlcNAc-derived intermediates will be required to determine whether enhanced nutrient utilization, mitigation of amino-sugar metabolic imbalance, or both underlie the protective effect of *NAG* overexpression. Likewise, the proposed contribution of chitin-derived GlcNAc recycling during late-stage growth remains to be tested directly. Further investigation of these mechanisms will help clarify how ZEA production influences primary nutrient metabolism and fungal physiology.

Collectively, our findings support a model in which ZEA production is coupled to an amino-sugar metabolic response in *F. graminearum*. Rather than functioning solely as a terminal product of secondary metabolism, ZEA appears to establish a physiological state that increases the cellular requirement for GlcNAc catabolism. Within this response, *NAG* serves as a critical metabolic node that supports fungal fitness under conditions of elevated ZEA production. Although the broader physiological and ecological significance of this connection remains to be established, its contribution may become particularly relevant under complex environmental conditions where fungi encounter fluctuating nutrient availability and interactions with other organisms. By uncovering a previously unrecognized connection between mycotoxin production and primary nutrient metabolism, our study provides a new perspective on the longstanding question of how fungal secondary metabolites can contribute to the physiology and fitness of their producers.

## Materials and Methods

### Fungal strains and cultivation conditions

The *Fusarium graminearum* wild-type strain (Z-3639) and the mutant strains generated in this study are listed in Supplementary Data 3. General culture media and standard laboratory protocols followed previously described methods ^44^. For genomic DNA isolation, fungal strains were cultivated in liquid complete medium (CM) for 3 days at 25 °C with shaking at 150 rpm, after which the mycelia were harvested via filtration. Conidiation was induced using carboxymethyl cellulose (CMC) liquid medium or yeast malt agar (YMA) medium ^45^. All strains, including the wild-type and its derivatives, were maintained as mycelial plugs in a 20% (v/v) glycerol solution at-80 °C for long-term storage.

### GlcNAc utilization assays

GlcNAc utilization was evaluated on carbon source-free minimal medium (MM−C), prepared by omitting the carbon source from the standard minimal-medium formulation. MM−C was supplemented with 1% (w/v) *N*-acetylglucosamine (GlcNAc), 1% (w/v) glucose, or both, as indicated for each experiment. GlcNAc concentrations were varied as specified in the corresponding figures.

Conditional *FgNAG* expression was evaluated using the *P_FCR1_-FgNAG* and *P_FCR1_-FgNAG*; *ZEB2LOE* strains grown on MM−C supplemented with 1% (w/v) glucose or GlcNAc. Conditional expression of *FgNAG* under the copper-responsive *FCR1* promoter was modulated by supplementation with 50 µM BCS or 25 µM CuSO₄. Colony morphology and radial growth were evaluated at 3 dpi.

Growth inhibition was calculated as [(radial growth of the reference strain or condition - radial growth of the test strain or condition)/radial growth of the reference strain or condition] × 100. Negative values indicate enhanced growth relative to the corresponding reference.

### Transcriptome analysis and functional enrichment

RNA-sequencing data generated in our previous study ^46^ were reanalyzed to investigate transcriptional responses associated with *ZEB2* overexpression. Differentially expressed genes (DEGs) between the wild-type and *ZEB2OE* strains were identified using a *P* value < 0.05 and a log₂ fold change ≥ 2. A total of 393 genes upregulated in the *ZEB2OE* strain were subjected to functional enrichment analysis using enrichment tools in FungiDB ^47^. Enriched terms and pathways with a *P* value < 0.05 were retained for further analysis. Enrichment results were visualized in R using the ggplot2 package, with dot size representing the number of genes and color indicating the *P* value.

### Identification of GlcNAc catabolic genes

Putative GlcNAc catabolic proteins in *F. graminearum* were identified by BLASTp searches using previously characterized proteins from *Neurospora crassa* as queries ^26^. Candidate proteins corresponding to the GlcNAc transporter Ngt, GlcNAc kinase Hxk, GlcNAc-6-phosphate deacetylase Nag, glucosamine-6-phosphate deaminase Dam, and the transcriptional regulator Ron were identified based on sequence similarity, genomic organization, and transcriptional responses observed in the *ZEB2OE* transcriptome. For most pathway components, the best-matching candidate was selected based on BLASTp sequence similarity and the available functional annotations. For Dam, candidate selection additionally considered its genomic proximity to the predicted *NAG*, *RON*, and *HXK* genes and its strong transcriptional induction in the *ZEB2OE* strain (Supplementary Data 2). Genomic organization and chromosomal locations of the candidate genes were examined using the *F. graminearum* genome annotation available in FungiDB.

### Nucleic acid manipulation and genetic modification

Genomic DNA was isolated from fungal mycelia using the cetyltrimethylammonium bromide (CTAB) method ^44^. Targeted gene deletion constructs were generated using the double-joint PCR (DJ-PCR) method in combination with a split-marker strategy ^48,49^. Approximately 1-kb 5′ and 3′ flanking regions of each target gene were amplified from wild-type genomic DNA and fused to fragments of a geneticin resistance cassette (*GEN*) by overlapping PCR. The resulting constructs were introduced into *F. graminearum* protoplasts by polyethylene glycol (PEG)-mediated transformation ^50^. Correct integration of the deletion constructs was verified by diagnostic PCR and Southern blot hybridization using the BioPrime™ DNA Labeling System and the North2South Chemiluminescent Hybridization and Detection Kit (Thermo Fisher Scientific, USA) (Supplementary Fig. 6).

For complementation and overexpression strain construction, the corresponding wild-type open reading frame was amplified and cloned using a yeast gap-repair approach ^51^. For complementation constructs, the coding region of the target gene was amplified together with approximately 1 kb of its native upstream promoter region. In contrast, the coding region was placed under the control of the constitutive *RP27* promoter for overexpression constructs. PCR-amplified inserts and XhoI-linearized pDL2 were co-transformed into *Saccharomyces cerevisiae* strain PJ69-4A for cloning. Recombinant plasmids were recovered using the Zymoprep Yeast Plasmid Miniprep Kit (Zymo Research, USA) and subsequently transformed into *Escherichia coli* DH10B. The resulting constructs were verified by sequencing (Bionics, Republic of Korea) and introduced into the indicated genetic backgrounds by fungal transformation.

For conditional regulation of *FgNAG*, the native *NAG* promoter was replaced with the copper-responsive *FCR1* promoter (*P_FCR1_-FgNAG*) as previously described ^31^. The *P_FCR1_*-containing fragment was amplified from the previously generated *P_FCR1_-FgIRE1* strain and assembled with the corresponding *NAG*-targeting fragments for integration immediately upstream of the *NAG* start codon. Transformation was performed using the same PEG-mediated protoplast transformation procedure described above, and correct integration was confirmed by diagnostic PCR and Southern blot analysis. Conditional strains were cultured in media supplemented with 25 µM CuSO₄ to repress *P_FCR1_* activity or with 50 µM BCS to relieve copper-mediated repression.

For construction of the *P_FCR1_-FgNAG*; *ZEB2LOE* strain, the *ZEB2L* overexpression construct was introduced into the *P_FCR1_-FgNAG* background, and elevated *ZEB2* expression was confirmed by RT-qPCR. All primers used for strain construction and verification are listed in Supplementary Data 4.

### Zearalenone quantification

Zearalenone (ZEA) extraction and quantification were conducted in accordance with a previously established protocol ^18,52^. Fungal strains were cultured in 20 mL of SG liquid medium at 25 °C for 7 days. Culture filtrates were separated from fungal mycelia by filtration through Whatman No. 2 filter paper and used for ZEA extraction. Culture filtrates were extracted with ethyl acetate, and the organic phase was collected and evaporated to dryness. The resulting residue was reconstituted in methanol/water (70:30, v/v) before HPLC analysis. ZEA was quantified using a Prominence high-performance liquid chromatography (HPLC) system (Shimadzu, Japan) equipped with a Shim-pack GIS C18 column (5 µm, 4.6 × 250 mm; Shimadzu). The mobile phase consisted of methanol/water (70:30, v/v) and was delivered isocratically at a flow rate of 1 mL/min. ZEA was detected by UV absorbance at 238 nm and identified and quantified by comparison with an authentic ZEA standard. A 10 mg/mL stock solution of authentic ZEA (Sigma-Aldrich, USA) was prepared in dimethyl sulfoxide (DMSO) and stored until use.

### RNA extraction and reverse transcription quantitative PCR (RT-qPCR)

For RNA isolation, fungal mycelia harvested under the indicated culture conditions were immediately frozen in liquid nitrogen and ground to a fine powder. Total RNA was extracted using the Easy-Spin Total RNA Extraction Kit (iNtRON Biotechnology, Republic of Korea) according to the manufacturer’s instructions. RNA quantity and purity were assessed using a NanoDrop 2000 spectrophotometer (Thermo Fisher Scientific).

First-strand cDNA was synthesized using SuperScript III Reverse Transcriptase (Thermo Fisher Scientific), according to the manufacturer’s instructions. Quantitative PCR was performed using iTaq Universal SYBR Green Supermix (Bio-Rad, USA) on a CFX Real-Time PCR Detection System (Bio-Rad). Relative transcript abundance was calculated using the 2^−ΔΔCt^ method, with endogenous cyclophilin gene (*CYP1*) used as the internal reference gene. Each experiment consisted of three or four independent biological replicates. Primer sequences are listed in Supplementary Data 4.

### ZEA and deoxynivalenol (DON) treatment assays

For analysis of ZEA-responsive gene expression, fungal strains were cultured in 10 mL of SG medium, and exogenous ZEA was added to a final concentration of 30 µM. Mycelia were harvested at 3 days post-inoculation (dpi), immediately frozen in liquid nitrogen, and subjected to RNA extraction and RT-qPCR as described above. Untreated cultures grown under the same conditions served as controls. For DON treatment, the wild-type strain was cultured in DON-inducing MMA medium ^53,54^ with or without 3 µM exogenous deoxynivalenol (DON) (Sigma-Aldrich). Mycelia were harvested at 3 and 7 dpi for RT-qPCR analysis.

### ZEA tolerance assay

For ZEA tolerance assays, the wild-type and two independent *NAG*-overexpressing strains were cultured on YMA medium to induce conidiation and inoculated into 96-well plates at a final concentration of 1 × 10⁴ conidia/mL in 0.5 × potato dextrose broth (PDB). Cultures were exposed to exogenous ZEA at concentrations ranging from 0 to 80 µM. Cellular metabolic activity was assessed at 2 dpi using PrestoBlue Cell Viability Reagent (Thermo Fisher Scientific) according to the manufacturer’s instructions ^55,56^. Fluorescence was measured using a microplate reader at an excitation wavelength of 570 nm and an emission wavelength of 600 nm. The inhibition rate was calculated as [(fluorescence signal of the untreated control − fluorescence signal of the ZEA-treated culture) / fluorescence signal of the untreated control] × 100.

## Statistical analysis

All experiments were performed using at least three independent biological replicates, unless otherwise indicated. Data are presented as the mean ± standard deviation. Comparisons between two groups were performed using Welch’s two-tailed *t*-test. For comparisons involving multiple strains or treatments, data were analyzed by one-way analysis of variance (ANOVA) followed by Tukey’s honestly significant difference (HSD) test. Differences were considered statistically significant at P < 0.05. Statistical analyses were performed using GraphPad Prism version 11.1.0 (GraphPad Software, USA) or R version 4.6.1. The statistical tests, sample sizes, and significance criteria used for individual experiments are indicated in the corresponding figure legends.

## Data Availability

The RNA-seq data used in this study are available in the NCBI Sequence Read Archive (SRA) under accession number PRJNA1521817.

## Acknowledgements

This study was supported by the Korea Institute of Planning and Evaluation for Technology in Food, Agriculture, and Forestry (IPET) through the Agriculture and Food Convergence Technologies Program for Research Manpower Development, funded by the Ministry of Agriculture, Food and Rural Affairs (MAFRA) (No. RS-2024-00398300) and the National Research Foundation of Korea (NRF) grants funded by the Korea government (MSIT) (No. RS-2025-00553624).

## Contributions

N.L., and H.S. contributed to conceptualization. N.L., J.W.H., and H.K. contributed to methodology. N.L. and J.W.H. performed the investigation. N.L., J.W.H., and J.P. contributed to validation. J.Y. and H.S. provided resources. N.L. prepared the visualization. N.L. and H.S. wrote the paper. H.S. supervised the study.

## Ethics declarations

**Ethics declaration**

Not applicable.

## References

1 Brakhage, A. A. Regulation of fungal secondary metabolism. Nat. Rev. Microbiol. 11, 21–32 (2013).

2 Keller, N. P. Fungal secondary metabolism: regulation, function and drug discovery. Nat. Rev. Microbiol. 17, 167–180 (2019).

3 Macheleidt, J. et al. Regulation and role of fungal secondary metabolites. Annu. Rev. Genet. 50, 371–392 (2016).

4 Spiteller, P. Chemical ecology of fungi. Nat. Prod. Rep. 32, 971–993 (2015).

5 Bills, G. F. & Gloer, J. B. Biologically active secondary metabolites from the fungi. Microbiol. Spectr. 4 (2016).

6 Raffa, N. & Keller, N. P. A call to arms: Mustering secondary metabolites for success and survival of an opportunistic pathogen. PLOS Pathog. 15, e1007606 (2019).

7 Alves de Castro, P., et al. *Aspergillus fumigatus* mitogen-activated protein kinase MpkA is involved in gliotoxin production and self-protection. Nat. Commun. 15, 33 (2024).

8 Schimmel, T. G., Coffman, A. D. & Parsons, S. J. Effect of butyrolactone I on the producing fungus, *Aspergillus terreus*. Appl. Environ. Microbiol. 64, 3707–3712 (1998).

9 Boone, C. H. T., Parker, K. A., Gutzmann, D. J., Atkin, A. L. & Nickerson, K. W. Farnesol as an antifungal agent: comparisons among MTLa and MTLα haploid and diploid *Candida albicans* and *Saccharomyces cerevisiae*. Front. Physiol. 14, 1207567 (2023).

10 Hall, R. A. et al. The quorum-sensing molecules farnesol/homoserine lactone and dodecanol operate via distinct modes of action in *Candida albicans*. Eukaryot. Cell 10, 1034–1042 (2011).

11 Hornby, J. M. et al. Quorum sensing in the dimorphic fungus *Candida albicans* is mediated by farnesol. Appl. Environ. Microbiol. 67, 2982–2992 (2001).

12 Finotti, E. et al. Aflatoxins are natural scavengers of reactive oxygen species. Sci. Rep. 11, 16024 (2021).

13 Zhen, H., Hu, Y., Xiong, K., Li, M. & Jin, W. The occurrence and biological control of zearalenone in cereals and cereal-based feedstuffs: a review. Food Addit. Contam. Part A. 41, 1344–1359 (2024).

14 Rai, A., Das, M. & Tripathi, A. Occurrence and toxicity of a fusarium mycotoxin, zearalenone. Crit. Rev. Food Sci. Nutr. 60, 2710–2729 (2020).

15 Caldwell, R. W., Tuite, J., Stob, M. & Baldwin, R. Zearalenone production by *Fusarium* species. Appl. Microbiol. 20, 31–34 (1970).

16 Kim, Y.-T. et al. Two different polyketide synthase genes are required for synthesis of zearalenone in *Gibberella zeae*. Mol. Microbiol. 58, 1102–1113 (2005).

17 Kim, J.-E., Son, H. & Lee, Y.-W. Biosynthetic mechanism and regulation of zearalenone in *Fusarium graminearum*. JSM Mycotoxins 68, 1–6 (2018).

18 Park, A. R. et al. Autoregulation of *ZEB2* expression for zearalenone production in *Fusarium graminearum*. Mol. Microbiol. 97, 942–956 (2015).

19 Tharanathan, R. N. & Kittur, F. S. Chitin—the undisputed biomolecule of great potential. Crit. Rev. Food Sci. Nutr. 43, 61–87 (2003).

20 Su, C., Lu, Y. & Liu, H. *N*-acetylglucosamine sensing by a *GCN5*-related *N*-acetyltransferase induces transcription via chromatin histone acetylation in fungi. Nat. Commun. 7, 12916 (2016).

21 Alvarez, F. J. & Konopka, J. B. Identification of an *N*-acetylglucosamine transporter that mediates hyphal induction in *Candida albicans*. Mol. Biol. Cell 18, 965–975 (2007).

22 Simonetti, N., Strippoli, V. & Cassone, A. Yeast-mycelial conversion induced by N-acetyl-D-glucosamine in *Candida albicans*. Nature 250, 344–346 (1974).

23 Camacho, E., et al. *N*-acetylglucosamine affects *Cryptococcus neoformans* cell-wall composition and melanin architecture. Microbiology 163, 1540–1556 (2017).

24 Bhatt, D. N., et al. *Magnaporthe oryzae* MoNdt80 is a transcriptional regulator of GlcNAc catabolic pathway involved in pathogenesis. Microbiol. Res. 239, 126550 (2020).

25 Gong, X. et al. Transcription factor RonA-driven GlcNAc catabolism is essential for growth, cell wall integrity, and pathogenicity in *Aspergillus fumigatus*. Microbiol. Spectr. 13 (2025).

26 Gaderer, R., Seidl-Seiboth, V., de Vries, R. P., Seiboth, B. & Kappel, L. *N*-acetylglucosamine, the building block of chitin, inhibits growth of *Neurospora crassa*. Fungal Genet. Biol. 107, 1–11 (2017).

27 Ullah, S. F., et al. *N*-acetylglucosamine sensing in the filamentous soil fungus *Trichoderma reesei*. FEBS J. 292, 3072–3090 (2025).

28 Kappel, L., Gaderer, R., Flipphi, M. & Seidl-Seiboth, V. The *N*-acetylglucosamine catabolic gene cluster in *Trichoderma reesei* is controlled by the Ndt80-like transcription factor RON1. Mol. Microbiol. 99, 640–657 (2016).

29 Kim, J.-E., et al. *Gibberella zeae* chitin synthase genes, GzCHS5 and GzCHS7, are required for hyphal growth, perithecia formation, and pathogenicity. Curr. Genet. 55, 449–459 (2009).

30 Son, H. et al. A novel transcription factor gene *FHS1* is involved in the DNA damage response in *Fusarium graminearum*. Sci. Rep. 6, 21572 (2016).

31 Kim, S. et al. Development of a versatile copper-responsive gene expression system in the plant-pathogenic fungus *Fusarium graminearum*. Mol. Plant Pathol. 22, 1427–1435 (2021).

32 Lee, N. et al. Membrane-associated compartmentalization of zearalenone biosynthetic enzymes and Syn2-associated zearalenone homeostasis in *Fusarium graminearum*. Microbiol. Res. 314, 128730 (2027).

33 Alves de Castro, P., et al. Regulation of gliotoxin biosynthesis and protection in *Aspergillus* species. PLOS Genet. 18, e1009965 (2022).

34 Kleijn, R. J. et al. Cytosolic NADPH metabolism in penicillin-G producing and non-producing chemostat cultures of *Penicillium chrysogenum*. Metab. Eng. 9, 112–123 (2007).

35 Panagiotou, G. et al. Studies of the production of fungal polyketides in *Aspergillus nidulans* by using systems biology tools. Appl. Environ. Microbiol. 75, 2212–2220 (2009).

36 Reyes, F., Calatayud, J., Vazquez, C. & Martínez, M. J. *β*-*N*-Acetylglucosaminidase from *Aspergillus nidulans* which degrades chitin oligomers during autolysis. FEMS Microbiol. Lett. 65, 83–87 (1989).

37 Perez-Leblic, M. I., Reyes, F., Martinez, M. J. & Lahoz, R. Cell wall degradation in the autolysis of filamentous fungi. Mycopathologia 80, 147–155 (1982).

38 Ellström, M. et al. The carbon starvation response of the ectomycorrhizal fungus *Paxillus involutus*. FEMS Microbiol. Ecol. 91, fiv027 (2015).

39 van Munster, J. M. et al. Systems approaches to predict the functions of glycoside hydrolases during the life cycle of *Aspergillus niger* using developmental mutants Δ*brlA* and Δ*flbA*. PLOS One 10, e0116269 (2015).

40 Josefsen, L. et al. Autophagy provides nutrients for nonassimilating fungal structures and is necessary for plant colonization but not for infection in the necrotrophic plant pathogen *Fusarium graminearum*. Autophagy 8, 326–337 (2012).

41 Kumar, M. J., Jamaluddin, M. S., Natarajan, K., Kaur, D. & Datta, A. The inducible N-acetylglucosamine catabolic pathway gene cluster in *Candida albicans*: Discrete N-acetylglucosamine-inducible factors interact at the promoter of *NAG1*. Proc. Natl. Acad. Sci. U. S. A. 97, 14218–14223 (2000).

42 Yamada-Okabe, T., Sakamori, Y., Mio, T. & Yamada-Okabe, H. Identification and characterization of the genes for *N*-acetylglucosamine kinase and *N*-acetylglucosamine-phosphate deacetylase in the pathogenic fungus *Candida albicans*. Eur. J. Biochem. 268, 2498–2505 (2001).

43 Naseem, S., Gunasekera, A., Araya, E. & Konopka, J. B. *N*-acetylglucosamine (GlcNAc) induction of hyphal morphogenesis and transcriptional responses in *Candida albicans* are not dependent on its metabolism. J. Biol. Chem. 286, 28671–28680 (2011).

44 Leslie, J. F. & Summerell, B. A. The Fusarium Laboratory Manual (Blackwell Publishing, Ames, IA, 2006).

45 Harris, S. D. Morphogenesis in germinating *Fusarium graminearum* macroconidia. Mycologia 97, 880–887 (2005).

46 Lee, N. et al. Transcriptome dissection of the Zeb2-associated network reveals a role for *NRPS15* in zearalenone biosynthesis in *Fusarium graminearum*. Preprint at bioRxiv 10.64898/2026.10.23.761285 (2026).

47 Basenko, E. Y. et al. What is new in FungiDB: a web-based bioinformatics platform for omics-scale data analysis for fungal and oomycete species. Genetics 227, iyae035 (2024).

48 Catlett, N. L., Lee, B.-N., Yoder, O. C. & Turgeon, B. G. Split-marker recombination for efficient targeted deletion of fungal genes. Fungal Genet. Rep. 50, 9–11 (2003).

49 Yu, J.-H. et al. Double-joint PCR: a PCR-based molecular tool for gene manipulations in filamentous fungi. Fungal Genet. Biol. 41, 973–981 (2004).

50 Han, Y.-K., Kim, M.-D., Lee, S.-H., Yun, S.-H. & Lee, Y.-W. A novel F-box protein involved in sexual development and pathogenesis in *Gibberella zeae*. Mol. Microbiol. 63, 768–779 (2007).

51 Zhou, X., Li, G. & Xu, J.-R. Efficient approaches for generating GFP fusion and epitope-tagging constructs in filamentous fungi. Methods Mol. Biol. 722, 199–212 (2011).

52 Bacon, C. W., Robbins, J. D. & Porter, J. K. Media for identification of *Gibberella zeae* and production of F-2-(Zearalenone). Appl. Environ. Microbiol. 33, 445–449 (1977).

53 Son, H., Lee, J., Park, A. R. & Lee, Y.-W. ATP citrate lyase is required for normal sexual and asexual development in *Gibberella zeae*. Fungal Genet. Biol. 48, 408–417 (2011).

54 Gardiner, D. M., Kazan, K. & Manners, J. M. Nutrient profiling reveals potent inducers of trichothecene biosynthesis in *Fusarium graminearum*. Fungal Genet. Biol. 46, 604–613 (2009).

55 Nguyen, H. T. et al. The Hsp90 inhibitor, monorden, is a promising lead compound for the development of novel fungicides. Front. Plant Sci. 11, 371 (2020).

56 Palomino, J.-C. et al. Resazurin microtiter assay plate: simple and inexpensive method for detection of drug resistance in *Mycobacterium tuberculosis*. Antimicrob. Agents Chemother. 46, 2720–2722 (2002).

