## Supplementary Information for "The mycotoxin zearalenone couples secondary metabolism to *N*-acetylglucosamine catabolism"

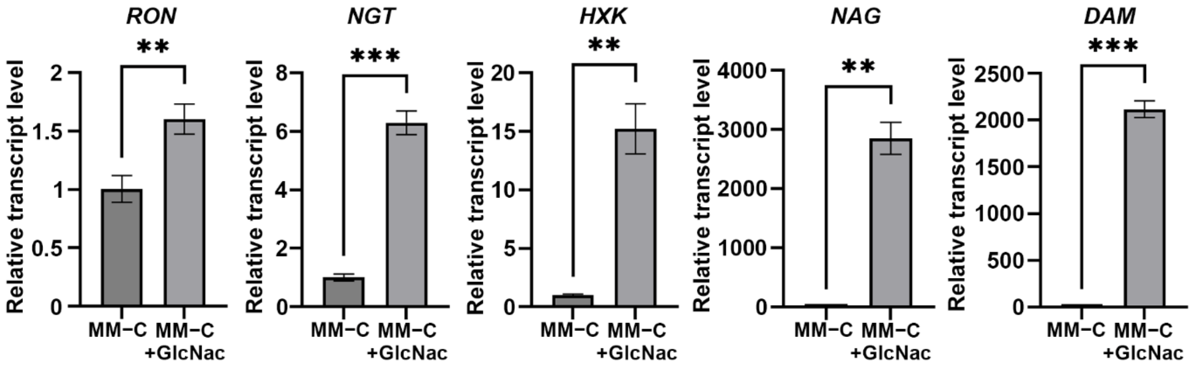


**Supplementary Fig. 1. GlcNAc induces expression of GlcNAc catabolic genes in *F. graminearum*.** Relative transcript levels of *RON*, *NGT*, *HXK*, *NAG*, and *DAM* were determined by RT-qPCR in the wild-type strain grown on MM−C with or without 1% (w/v) GlcNAc supplementation. Transcript levels were normalized to *CYP1* and are presented relative to those under the MM−C condition. Data are presented as mean ± SD from four independent biological replicates. Statistical significance was determined using Welch's two tailed *t*-test. **$P < 0.01$; ***$P < 0.001$.

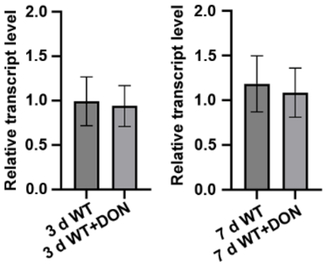


**Supplementary Fig. 2. Exogenous DON does not alter *NAG* expression in *F. graminearum*.** Relative *NAG* transcript levels were determined by RT-qPCR in the wild-type strain cultured under DON-inducing conditions with or without 3 μM exogenous deoxynivalenol (DON) for 3 or 7 days. Transcript levels were normalized to *CYP1* and are presented relative to the corresponding untreated wild-type control at each time point. Data are presented as mean ± SD from three independent biological replicates. No significant differences were detected between untreated and DON-treated samples by Welch's two-tailed *t*-test.

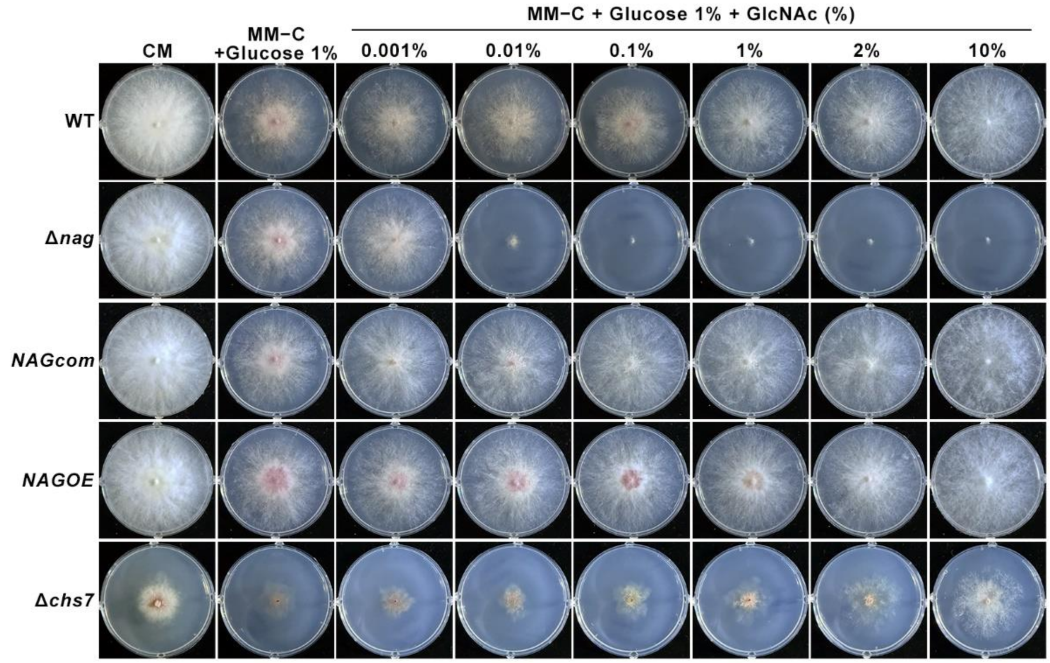


**Supplementary Fig. 3. The growth defect of Δ*nag* is GlcNAc-dependent and restored by *NAG* complementation or overexpression.** Representative colony morphology of the wild-type, Δ*nag*, *NAGcom*, *NAGOE*, and Δ*chs7* strains grown on complete medium (CM), carbon source-free minimal medium supplemented with 1% (w/v) glucose, or MM−C + 1% (w/v) glucose supplemented with increasing concentrations of GlcNAc 0.001–10% (w/v). The Δ*nag* mutant exhibited a pronounced concentration-dependent growth defect in response to GlcNAc, whereas growth was restored by *NAG* complementation or overexpression. In contrast, Δ*chs7* showed progressively enhanced growth with increasing GlcNAc concentrations.

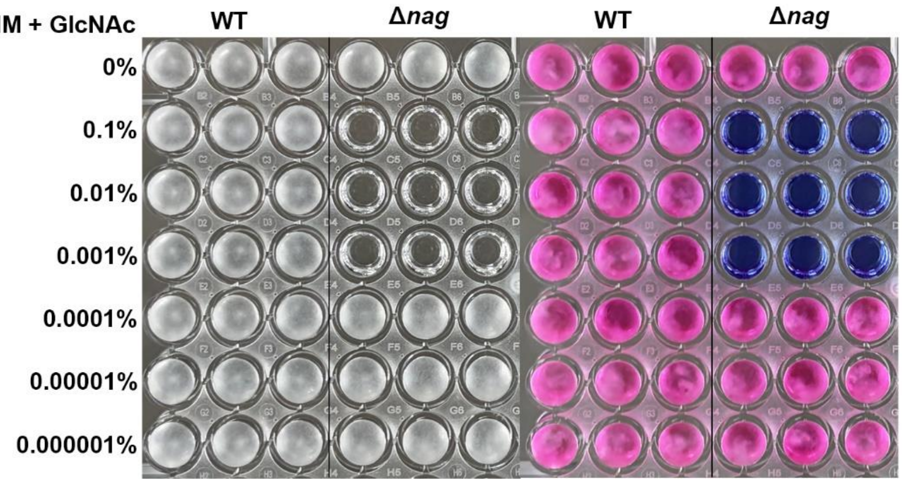


**Supplementary Fig. 4. GlcNAc causes a concentration-dependent reduction in growth and metabolic activity of the Δ*nag* mutant.** The wild-type and Δ*nag* strains were cultured in MM supplemented with the indicated concentrations of GlcNAc 0–0.1% (w/v). Representative cultures are shown on the left, and PrestoBlue reactions reflecting cellular metabolic activity are shown on the right. A color change of PrestoBlue from blue to pink indicates increased cellular metabolic activity.

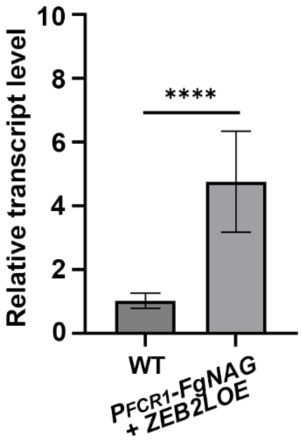


**Supplementary Fig. 5. Confirmation of *ZEB2L* overexpression in the *$P_{FCR1}$-FgNAG*; *ZEB2LOE* strain.** Relative *ZEB2* transcript levels were determined by RT-qPCR in the wild-type and *$P_{FCR1}$-FgNAG*; *ZEB2LOE* strains. Transcript levels were normalized to *CYP1* and are presented relative to the wild type. Data are presented as the mean ± SD from three independent biological replicates. Statistical significance was determined using Welch's two-tailed *t*-test. ****$P < 0.0001$.

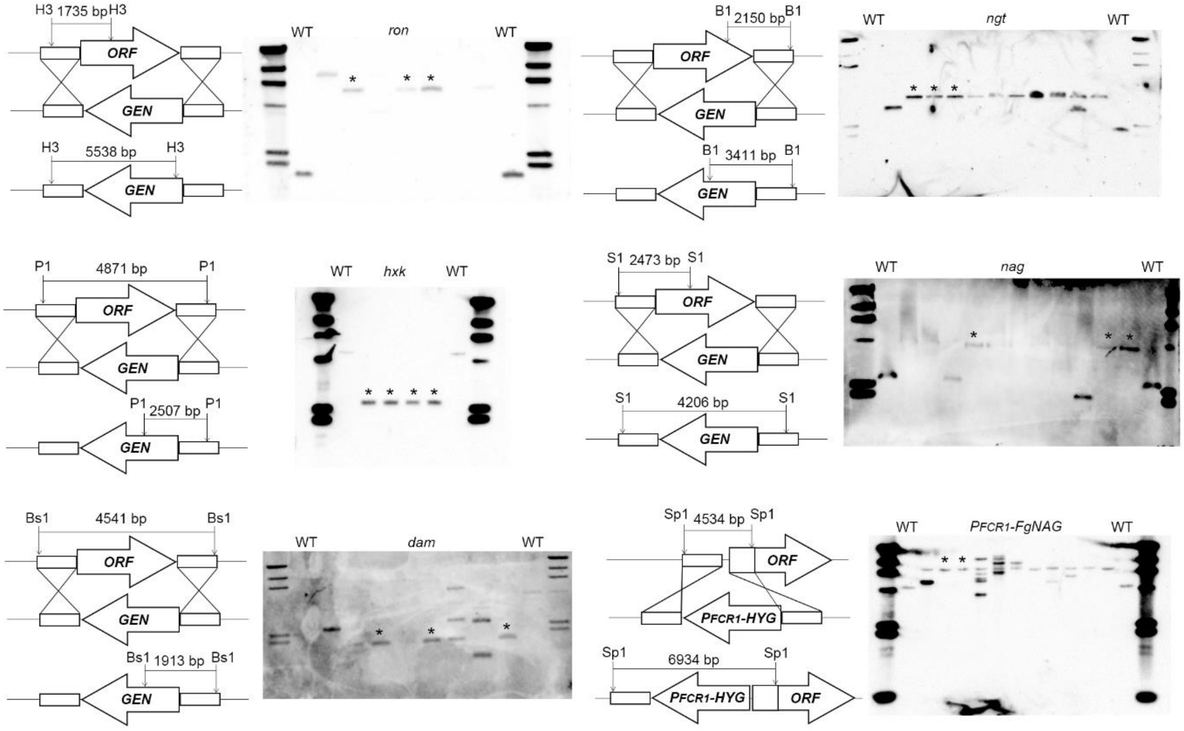


**Supplementary Fig. 6. Southern blot verification of the GlcNAc catabolic gene mutants and the conditional *NAG* expression strain.** Schematic representations of the targeted genetic modifications and Southern blot analyses are shown for knockout mutants and $P_{FCR1}$-*FgNAG*. Genomic DNA from the wild-type (WT) and the indicated mutant strains was digested with the restriction enzymes shown in each schematic and subjected to Southern blot analysis using the corresponding gene-specific probes. Uncropped Southern blot images are presented. Asterisks indicate transformants selected to establish the corresponding verified strains. B1, BsaI; S1, SacI; P1, PstI; H3, HindIII; BS1, BsmI; Sp1, SpeI; *GEN*, geneticin resistance cassette; $P_{FCR1}$-*HYG*, *FCR1* promoter carrying hygromycin B resistance cassette; WT, wild-type strain Z-3639.

**Supplementary Data 1. Genes upregulated in the *ZEB2OE* strain relative to the wild type.**

| Gene ID | Description | Log2(FC) | P-adj |
|---|---|---|---|
| FGRAMPH1_01G15627 | ATP binding cassette transporter | 15.62344671 | 0 |
| FGRAMPH1_01G05755 | hypothetical protein | 14.40075873 | 0 |
| FGRAMPH1_01G05747 | K+ channel | 13.4777521 | 1.6504E-214 |
| FGRAMPH1_01G05753 | hypothetical protein | 12.9502218 | 1.8704E-152 |
| FGRAMPH1_01G05745 | nonribosomal peptide synthetase | 12.66883079 | 8.1201E-162 |
| FGRAMPH1_01G15629 | transcription factor | 12.5552965 | 3.3815E-160 |
| FGRAMPH1_01G05757 | serine/threonine-protein kinase 6 | 10.37289665 | 1.1095E-154 |
| FGRAMPH1_01G05751 | hypothetical protein | 8.661256369 | 3.73732E-36 |
| FGRAMPH1_01G05337 | 2`-hydroxyisoflavone reductase | 8.574991875 | 9.4574E-203 |
| FGRAMPH1_01G05809 | hypothetical protein | 8.414717326 | 1.558E-239 |
| FGRAMPH1_01G13921 | hypothetical protein | 8.334175629 | 4.24198E-56 |
| FGRAMPH1_01G05749 | hypothetical protein | 7.251701872 | 1.19897E-22 |
| FGRAMPH1_01G00153 | hypothetical protein | 6.724865865 | 6.9196E-39 |
| FGRAMPH1_01G13695 | hypothetical protein | 6.060946194 | 5.8228E-106 |
| FGRAMPH1_01G20433 | isoflavone reductase family | 6.0018514 | 3.5536E-120 |
| FGRAMPH1_01G18451 | Bifunctional xylanase deacetylase | 5.965076036 | 3.75983E-82 |
| FGRAMPH1_01G12517 | transcription factor | 5.947906011 | 1.1065E-216 |
| FGRAMPH1_01G12233 | nadph2:quinone reductase | 5.914123716 | 1.16316E-82 |
| FGRAMPH1_01G12719 | hypothetical protein | 5.868644129 | 3.82786E-38 |
| FGRAMPH1_01G13655 | hypothetical protein | 5.79368526 | 1.40361E-43 |
| FGRAMPH1_01G22509 | hypothetical protein | 5.705603717 | 3.4319E-32 |
| FGRAMPH1_01G01377 | gnat family acetyltransferase | 5.368936298 | 3.41162E-24 |
| FGRAMPH1_01G12813 | hypothetical protein | 5.227264752 | 3.0882E-110 |
| FGRAMPH1_01G00541 | related to integral membrane PTH11 | 5.12731084 | 2.4049E-146 |
| FGRAMPH1_01G21205 | hypothetical protein | 5.088564106 | 5.33246E-37 |
| FGRAMPH1_01G16831 | potassium sodium efflux p-type fungal-type | 5.00357194 | 4.14729E-90 |
| FGRAMPH1_01G15885 | aldo-keto reductase yakc | 4.976565004 | 4.31725E-47 |
| FGRAMPH1_01G09231 | hypothetical protein | 4.867317004 | 5.97087E-11 |
| FGRAMPH1_01G09851 | n-acetylglucosamine-6-phosphate deacetylase | 4.734823396 | 1.22031E-60 |
| FGRAMPH1_01G25247 | d-lactate dehydrogenase | 4.717000255 | 4.12477E-19 |
| FGRAMPH1_01G12417 | glycerol kinase | 4.618492207 | 5.48211E-29 |
| FGRAMPH1_01G10141 | gluconate 5-dehydrogenase | 4.609273825 | 1.25307E-50 |
| FGRAMPH1_01G11959 | hypothetical protein | 4.584861871 | 1.09813E-11 |
| FGRAMPH1_01G12215 | unnamed protein product | 4.537429833 | 1.94319E-64 |
| FGRAMPH1_01G25935 | hypothetical protein | 4.504652517 | 1.32109E-65 |
| FGRAMPH1_01G00467 | GTP-binding RHO3 | 4.429982379 | 1.35185E-45 |
| FGRAMPH1_01G13977 | amidohydrolase family | 4.34981591 | 7.41844E-10 |
| FGRAMPH1_01G00181 | hypothetical protein | 4.342408301 | 6.05741E-10 |
| FGRAMPH1_01G15631 | glutamyl-trna synthetase | 4.249497819 | 3.94727E-83 |

| | | | |
|---|---|---|---|
| FGRAMPH1_01G15351 | hypothetical protein | 4.204055219 | 4.6709E-14 |
| FGRAMPH1_01G13401 | Nitrosoguanidine resistance SNG1 | 4.162537576 | 6.34465E-48 |
| FGRAMPH1_01G09059 | hypothetical protein | 4.061508878 | 1.1129E-08 |
| FGRAMPH1_01G03311 | alternative oxidase | 4.045083195 | 3.75527E-27 |
| FGRAMPH1_01G11927 | hypothetical protein | 4.04154512 | 1.4741E-18 |
| FGRAMPH1_01G11525 | sulfite oxidase | 3.997263433 | 1.4758E-44 |
| FGRAMPH1_01G17423 | cell wall manno | 3.815678456 | 1.44412E-30 |
| FGRAMPH1_01G16231 | hypothetical protein | 3.809338316 | 9.96738E-19 |
| FGRAMPH1_01G09373 | nonribosomal peptide synthetase | 3.805652205 | 1.1576E-17 |
| FGRAMPH1_01G12519 | oxidoreductase yulF | 3.793950779 | 1.5507E-57 |
| FGRAMPH1_01G09857 | glucosamine-6-phosphate isomerase | 3.775410358 | 1.1251E-115 |
| FGRAMPH1_01G21625 | Endo-1,4-beta-xylanase B | 3.754387741 | 3.4621E-23 |
| FGRAMPH1_01G16245 | retinol dehydrogenase | 3.754196746 | 1.76186E-16 |
| FGRAMPH1_01G14779 | hypothetical protein | 3.719913639 | 1.62324E-47 |
| FGRAMPH1_01G25277 | Pentalenolactone D synthase | 3.715598778 | 1.06354E-18 |
| FGRAMPH1_01G02873 | 30 kDa heat shock protein | 3.698912585 | 4.11416E-97 |
| FGRAMPH1_01G05743 | Heterokaryon incompatibility OR allele | 3.679345679 | 5.72309E-19 |
| FGRAMPH1_01G20819 | hypothetical protein | 3.671454587 | 5.05308E-14 |
| FGRAMPH1_01G08115 | hypothetical protein | 3.63550599 | 5.31879E-73 |
| FGRAMPH1_01G16001 | cycloheximide-inducible CIP70 (cytochrome P450 family) | 3.628938579 | 1.2396E-05 |
| FGRAMPH1_01G26655 | hypothetical protein | 3.62839636 | 3.85818E-27 |
| FGRAMPH1_01G10937 | ATPase | 3.620938295 | 9.91334E-50 |
| FGRAMPH1_01G09147 | unnamed protein product | 3.585997 | 1.71026E-06 |
| FGRAMPH1_01G01381 | leptomycin b resistance pmd1 | 3.559023324 | 8.29807E-64 |
| FGRAMPH1_01G13079 | hypothetical protein | 3.52436965 | 1.43807E-17 |
| FGRAMPH1_01G09061 | hypothetical protein | 3.494474234 | 1.44929E-07 |
| FGRAMPH1_01G12569 | related to oxidoreductase | 3.489041984 | 3.35722E-60 |
| FGRAMPH1_01G02875 | hypothetical protein FOWG_07428 | 3.488737986 | 2.57934E-34 |
| FGRAMPH1_01G05341 | lipoxygenase 1 | 3.470464114 | 6.99233E-12 |
| FGRAMPH1_01G08411 | hypothetical protein | 3.460551177 | 1.95198E-05 |
| FGRAMPH1_01G11785 | hypothetical protein | 3.449834302 | 4.54546E-22 |
| FGRAMPH1_01G21053 | nonribosomal peptide synthetase | 3.429067319 | 1.51819E-28 |
| FGRAMPH1_01G03065 | hypothetical protein | 3.398241173 | 1.16638E-28 |
| FGRAMPH1_01G17149 | unnamed protein product | 3.395360844 | 6.47562E-05 |
| FGRAMPH1_01G11193 | hypothetical protein | 3.376944713 | 2.91226E-08 |
| FGRAMPH1_01G16595 | unnamed protein product | 3.373490499 | 1.1761E-23 |
| FGRAMPH1_01G14737 | unspecified product | 3.372639726 | 2.10004E-53 |
| FGRAMPH1_01G13739 | glutathione s-transferase p 10 | 3.354773991 | 7.80465E-79 |
| FGRAMPH1_01G12755 | 30 kda heat shock | 3.348944883 | 5.14111E-41 |
| FGRAMPH1_01G11527 | catalase | 3.331513561 | 1.32188E-06 |
| FGRAMPH1_01G26653 | hypothetical protein | 3.328534295 | 1.6106E-23 |

| | | | |
|---|---|---|---|
| FGRAMPH1_01G05093 | cytochrome p450 3a7 | 3.324615353 | 9.40663E-11 |
| FGRAMPH1_01G04389 | L-lactate dehydrogenase (cytochrome) | 3.32426811 | 3.94349E-08 |
| FGRAMPH1_01G12869 | endothiapepsin precursor | 3.303283884 | 1.5645E-103 |
| FGRAMPH1_01G04773 | aspartyl-trna synthetase | 3.297906783 | 1.03337E-50 |
| FGRAMPH1_01G16407 | indoleamine -dioxygenase | 3.297873232 | 2.32616E-44 |
| FGRAMPH1_01G28067 | hypothetical protein | 3.269948557 | 1.08822E-06 |
| FGRAMPH1_01G12217 | hypothetical protein | 3.239195295 | 2.93954E-12 |
| FGRAMPH1_01G11499 | Tip elongation aberrant 3 | 3.223867956 | 9.76779E-13 |
| FGRAMPH1_01G21921 | hypothetical protein | 3.214149625 | 2.55342E-09 |
| FGRAMPH1_01G13995 | Cut9-interacting scn1 | 3.21130338 | 5.52616E-09 |
| FGRAMPH1_01G15637 | glycoside hydrolase family 76 | 3.204217868 | 5.03541E-17 |
| FGRAMPH1_01G03717 | btb poz containing | 3.197524095 | 2.68487E-13 |
| FGRAMPH1_01G15523 | unnamed protein product | 3.193325225 | 3.2363E-09 |
| FGRAMPH1_01G11829 | BAHD acyltransferase DCR | 3.185422862 | 6.19227E-09 |
| FGRAMPH1_01G28015 | 2,3-dihydroxybenzoic acid decarboxylase | 3.170923074 | 7.15058E-07 |
| FGRAMPH1_01G07889 | egg shell | 3.167823007 | 2.06731E-28 |
| FGRAMPH1_01G21265 | hypothetical protein | 3.161774039 | 1.63717E-05 |
| FGRAMPH1_01G14529 | NAD-specific glutamate dehydrogenase | 3.160124415 | 1.47678E-39 |
| FGRAMPH1_01G19711 | hypothetical protein | 3.149184536 | 4.10479E-30 |
| FGRAMPH1_01G15357 | hypothetical protein | 3.136927442 | 1.58219E-06 |
| FGRAMPH1_01G20813 | ww oxidoreductase | 3.135223172 | 3.85445E-18 |
| FGRAMPH1_01G25655 | Repressible acid phosphatase | 3.126815213 | 2.23406E-60 |
| FGRAMPH1_01G16537 | hypothetical protein | 3.121320345 | 1.13348E-10 |
| FGRAMPH1_01G13069 | related to neutral amino acid permease | 3.114337772 | 9.38529E-12 |
| FGRAMPH1_01G18017 | heat shock protein HSP98 | 3.111087406 | 2.98993E-70 |
| FGRAMPH1_01G21057 | abc transporter | 3.101760453 | 1.71912E-15 |
| FGRAMPH1_01G26949 | hypothetical protein | 3.096778756 | 5.67716E-26 |
| FGRAMPH1_01G05099 | unnamed protein product | 3.070961379 | 1.19855E-07 |
| FGRAMPH1_01G10315 | hypothetical protein | 3.069928582 | 2.55686E-05 |
| FGRAMPH1_01G25135 | 3-dehydroquinate synthase | 3.065660455 | 3.17411E-07 |
| FGRAMPH1_01G14927 | Zinc finger rsv2 | 3.050103987 | 2.49123E-31 |
| FGRAMPH1_01G12419 | aquaglyceroporin like other eukaryote | 3.046142838 | 1.8995E-106 |
| FGRAMPH1_01G26135 | hypothetical protein | 3.044299266 | 6.87297E-21 |
| FGRAMPH1_01G14993 | mfs multidrug | 3.037611003 | 3.6868E-25 |
| FGRAMPH1_01G28065 | N amino acid transport system protein | 3.035912972 | 4.67538E-41 |
| FGRAMPH1_01G15353 | hypothetical protein | 3.029457341 | 0.000499143 |
| FGRAMPH1_01G21291 | cyanamide hydratase | 3.0187732 | 1.2704E-13 |
| FGRAMPH1_01G26491 | hypothetical protein | 3.014231608 | 4.87004E-28 |
| FGRAMPH1_01G02125 | integral membrane | 3.013856026 | 1.12989E-38 |
| FGRAMPH1_01G13125 | chitin deacetylase | 3.013354457 | 7.40241E-29 |
| FGRAMPH1_01G13991 | hypothetical protein | 3.007852623 | 0.000316441 |
| FGRAMPH1_01G09669 | hypothetical protein | 3.003658395 | 1.36288E-52 |

| | | | |
|---|---|---|---|
| FGRAMPH1_01G17019 | lipase rog1 | 2.991331893 | 2.66895E-36 |
| FGRAMPH1_01G13767 | epoxide hydrolase | 2.986482731 | 6.39722E-37 |
| FGRAMPH1_01G28291 | hypothetical protein | 2.984883758 | 2.60916E-18 |
| FGRAMPH1_01G04105 | alcohol dehydrogenase (NADP+) | 2.981289744 | 1.68068E-10 |
| FGRAMPH1_01G12063 | hypothetical protein | 2.974071234 | 7.66763E-46 |
| FGRAMPH1_01G25327 | integral membrane | 2.967332431 | 5.96255E-31 |
| FGRAMPH1_01G02553 | related to thermoresistant gluconokinase | 2.961355663 | 4.31935E-48 |
| FGRAMPH1_01G24373 | NADH dehydrogenase | 2.96134169 | 1.9001E-47 |
| FGRAMPH1_01G06471 | WD repeat-containing | 2.956741675 | 6.86131E-67 |
| FGRAMPH1_01G27629 | hypothetical protein | 2.955963371 | 5.94711E-56 |
| FGRAMPH1_01G15355 | hypothetical protein | 2.938650948 | 1.2451E-17 |
| FGRAMPH1_01G21529 | adam 8 precursor | 2.913183942 | 0.000781117 |
| FGRAMPH1_01G15717 | hypothetical protein | 2.906861721 | 5.42123E-08 |
| FGRAMPH1_01G26017 | hypothetical protein | 2.904211472 | 3.84457E-09 |
| FGRAMPH1_01G25185 | histone acetyltransferase hpa2 | 2.8910289 | 9.55341E-05 |
| FGRAMPH1_01G12825 | zinc-type alcohol dehydrogenase | 2.889059013 | 5.26289E-07 |
| FGRAMPH1_01G20115 | hypothetical protein | 2.839667525 | 4.16315E-15 |
| FGRAMPH1_01G15607 | branched-chain amino acid aminotransferase | 2.835957157 | 1.71661E-56 |
| FGRAMPH1_01G18837 | glutamine amidotransferase | 2.834382569 | 8.69288E-44 |
| FGRAMPH1_01G08117 | hypothetical protein | 2.815242479 | 4.8974E-20 |
| FGRAMPH1_01G27627 | hypothetical protein | 2.815052378 | 2.90266E-07 |
| FGRAMPH1_01G15633 | small secreted | 2.81135272 | 4.28947E-29 |
| FGRAMPH1_01G08513 | hypothetical protein | 2.801194825 | 3.04053E-05 |
| FGRAMPH1_01G11725 | hypothetical protein | 2.798257839 | 2.40058E-31 |
| FGRAMPH1_01G13693 | 6-hydroxy-d-nicotine oxidase | 2.793021193 | 2.51468E-14 |
| FGRAMPH1_01G00889 | hypothetical protein | 2.785995373 | 8.61455E-22 |
| FGRAMPH1_01G23929 | hypothetical protein | 2.78587639 | 3.9833E-13 |
| FGRAMPH1_01G09229 | NAD-dependent dehydratase | 2.775862546 | 0.000877046 |
| FGRAMPH1_01G14253 | NADH oxidase | 2.775714703 | 8.08276E-42 |
| FGRAMPH1_01G00405 | Zinc-type alcohol dehydrogenase | 2.765195511 | 1.16537E-34 |
| FGRAMPH1_01G08369 | hypothetical protein | 2.762016056 | 2.50682E-28 |
| FGRAMPH1_01G27007 | hypothetical protein | 2.76186776 | 1.49723E-08 |
| FGRAMPH1_01G27087 | hypothetical protein | 2.756804323 | 3.25324E-09 |
| FGRAMPH1_01G20545 | mip family channel | 2.749147173 | 0.00017412 |
| FGRAMPH1_01G04309 | 2`-hydroxyisoflavone reductase | 2.746253076 | 0.000171548 |
| FGRAMPH1_01G03089 | hypothetical protein | 2.739763499 | 0.000203871 |
| FGRAMPH1_01G13413 | hypothetical protein | 2.736400963 | 3.03178E-32 |
| FGRAMPH1_01G22001 | related to acyl- transferases carnitine dehydratase | 2.733812387 | 8.71425E-10 |
| FGRAMPH1_01G12597 | hypothetical protein | 2.73244493 | 6.45668E-21 |
| FGRAMPH1_01G23997 | hypothetical protein | 2.729416789 | 2.87851E-29 |
| FGRAMPH1_01G17823 | glycoside hydrolase family 16 partial | 2.729241177 | 6.91352E-21 |
| FGRAMPH1_01G24155 | hypothetical protein | 2.717170234 | 8.41016E-08 |

| | | | |
|---|---|---|---|
| FGRAMPH1_01G26769 | hypothetical protein | 2.716214712 | 2.89733E-39 |
| FGRAMPH1_01G00885 | aureobasidin resistance aur1 | 2.712409497 | 4.81066E-51 |
| FGRAMPH1_01G16399 | hypothetical protein | 2.710264917 | 1.98069E-25 |
| FGRAMPH1_01G19569 | hypothetical protein | 2.704918902 | 6.15763E-14 |
| FGRAMPH1_01G18993 | hypothetical protein | 2.701679098 | 4.68491E-11 |
| FGRAMPH1_01G26379 | atp-dependent rna helicase dbp7 | 2.69506641 | 1.58277E-13 |
| FGRAMPH1_01G13755 | esterase lipase | 2.694071947 | 1.22889E-09 |
| FGRAMPH1_01G03981 | integral membrane | 2.687011293 | 2.5282E-14 |
| FGRAMPH1_01G00183 | 3-hydroxybenzoate 6-hydroxylase 1 | 2.685955022 | 6.21368E-16 |
| FGRAMPH1_01G16611 | hypothetical protein | 2.684671659 | 2.23872E-17 |
| FGRAMPH1_01G00441 | high affinity methionine permease | 2.671067225 | 6.5537E-09 |
| FGRAMPH1_01G12681 | NADPH2 dehydrogenase chain OYE2 | 2.66746172 | 1.44948E-21 |
| FGRAMPH1_01G07531 | an1-type zinc finger | 2.666944769 | 1.90252E-42 |
| FGRAMPH1_01G17673 | hypothetical protein | 2.66338834 | 4.61644E-08 |
| FGRAMPH1_01G16933 | pyridoxine biosynthesis protein PDX1 | 2.656324012 | 1.75749E-14 |
| FGRAMPH1_01G13613 | unnamed protein product | 2.650312893 | 1.01376E-23 |
| FGRAMPH1_01G20965 | 2-isopropylmalate synthase | 2.648964925 | 9.23134E-12 |
| FGRAMPH1_01G25571 | hypothetical protein | 2.64773065 | 1.63419E-43 |
| FGRAMPH1_01G03447 | minor allergen alt a 7 | 2.64748861 | 3.79377E-55 |
| FGRAMPH1_01G14075 | amidase | 2.645133574 | 1.43984E-40 |
| FGRAMPH1_01G01175 | hypothetical protein FPSE_02030 | 2.64451376 | 2.33476E-46 |
| FGRAMPH1_01G15841 | hypothetical protein | 2.643538066 | 9.22704E-11 |
| FGRAMPH1_01G10123 | hypothetical protein | 2.643188248 | 1.44297E-29 |
| FGRAMPH1_01G16151 | Ent-kaurene oxidase | 2.642924245 | 5.56917E-13 |
| FGRAMPH1_01G24789 | hypothetical protein | 2.636066582 | 2.6894E-05 |
| FGRAMPH1_01G05839 | s- glutathione dehydrogenase alcohol dehydrogenase | 2.622665572 | 0.000229171 |
| FGRAMPH1_01G24787 | hypothetical protein | 2.61112421 | 2.00824E-36 |
| FGRAMPH1_01G17553 | hypothetical protein | 2.610264372 | 9.95997E-12 |
| FGRAMPH1_01G04203 | hypothetical protein | 2.601962961 | 3.14993E-19 |
| FGRAMPH1_01G03569 | Latex clearing | 2.597757745 | 2.31419E-12 |
| FGRAMPH1_01G18269 | hypothetical protein | 2.596918614 | 2.11157E-28 |
| FGRAMPH1_01G06041 | hypothetical protein | 2.573427832 | 5.53091E-22 |
| FGRAMPH1_01G06601 | endothelin-converting enzyme 2 | 2.568708282 | 3.28666E-10 |
| FGRAMPH1_01G25329 | hypothetical protein | 2.568085591 | 1.51742E-05 |
| FGRAMPH1_01G01153 | hypothetical protein | 2.567667766 | 9.21304E-09 |
| FGRAMPH1_01G16241 | nadph dehydrogenase | 2.559752065 | 4.41071E-23 |
| FGRAMPH1_01G13131 | alcohol dehydrogenase | 2.556573129 | 2.56679E-08 |
| FGRAMPH1_01G11179 | hypothetical protein | 2.555280658 | 4.11557E-07 |
| FGRAMPH1_01G19571 | hypothetical protein | 2.550285697 | 7.58537E-17 |
| FGRAMPH1_01G05105 | hypothetical protein | 2.545274219 | 1.22683E-14 |
| FGRAMPH1_01G06439 | related to tetracycline resistance s | 2.543212977 | 4.45995E-43 |
| FGRAMPH1_01G10131 | hypothetical protein | 2.541671329 | 1.63336E-33 |

| | | | |
|---|---|---|---|
| FGRAMPH1_01G13779 | WW domain-containing oxidoreductase | 2.540840129 | 1.947E-07 |
| FGRAMPH1_01G10661 | hypothetical protein | 2.534333047 | 2.61639E-09 |
| FGRAMPH1_01G01375 | siderophore iron transporter mirB | 2.533400873 | 3.41162E-24 |
| FGRAMPH1_01G11827 | hypothetical protein | 2.529292622 | 1.20376E-05 |
| FGRAMPH1_01G01235 | integral membrane | 2.52225357 | 7.70906E-41 |
| FGRAMPH1_01G07983 | hypothetical protein | 2.520050222 | 1.43046E-06 |
| FGRAMPH1_01G19115 | phospholipase a2 | 2.519992793 | 2.35957E-23 |
| FGRAMPH1_01G02619 | hypothetical protein | 2.508877911 | 2.90973E-07 |
| FGRAMPH1_01G19709 | hypothetical protein | 2.50313328 | 3.1064E-08 |
| FGRAMPH1_01G01217 | hypothetical protein | 2.500274671 | 5.50518E-10 |
| FGRAMPH1_01G17541 | hypothetical protein | 2.495248064 | 1.63608E-08 |
| FGRAMPH1_01G13757 | General amino-acid permease GAP1 | 2.494146363 | 9.32014E-37 |
| FGRAMPH1_01G24705 | hypothetical protein | 2.485441143 | 1.82549E-13 |
| FGRAMPH1_01G27453 | pectate lyase B precursor | 2.47392053 | 5.50773E-10 |
| FGRAMPH1_01G13625 | major facilitator superfamily transporter | 2.468702933 | 3.53176E-40 |
| FGRAMPH1_01G23765 | hypothetical protein | 2.465883733 | 4.42753E-13 |
| FGRAMPH1_01G08121 | phosphoenolpyruvate synthase | 2.462864196 | 1.85676E-21 |
| FGRAMPH1_01G26679 | hypothetical protein | 2.45269444 | 5.22245E-06 |
| FGRAMPH1_01G21537 | GMC oxidoreductase | 2.451690087 | 1.61078E-40 |
| FGRAMPH1_01G08441 | 3-dehydroshikimate dehydratase | 2.448142888 | 1.28936E-13 |
| FGRAMPH1_01G04211 | related to transcription SIN3 binding | 2.427594851 | 1.22641E-17 |
| FGRAMPH1_01G19201 | mechanosensitive ion channel family | 2.427025127 | 1.08323E-59 |
| FGRAMPH1_01G27119 | hypothetical protein | 2.425804813 | 1.43937E-40 |
| FGRAMPH1_01G10339 | phospholipid-translocating atpase | 2.425247804 | 1.89557E-22 |
| FGRAMPH1_01G13169 | sh3 ysc84 1 | 2.419458195 | 1.65412E-16 |
| FGRAMPH1_01G10323 | pirin (iron-binding nuclear ) | 2.414097274 | 3.60694E-42 |
| FGRAMPH1_01G12327 | hypothetical protein | 2.40838049 | 3.08328E-24 |
| FGRAMPH1_01G25955 | aldo keto reductase | 2.405769866 | 9.36867E-37 |
| FGRAMPH1_01G24543 | hypothetical protein | 2.404621288 | 5.42073E-27 |
| FGRAMPH1_01G13339 | sensor hoxx | 2.403775165 | 9.4682E-15 |
| FGRAMPH1_01G24739 | hypothetical protein | 2.40133009 | 1.4983E-17 |
| FGRAMPH1_01G25613 | hypothetical protein | 2.396285808 | 3.2762E-17 |
| FGRAMPH1_01G10501 | hypothetical protein FPSE_11556 | 2.390655801 | 1.24593E-12 |
| FGRAMPH1_01G19567 | hypothetical protein | 2.390283385 | 0.000206609 |
| FGRAMPH1_01G01471 | general stress protein 39 | 2.386836031 | 2.96624E-14 |
| FGRAMPH1_01G07575 | hypothetical protein | 2.386188584 | 2.1673E-15 |
| FGRAMPH1_01G15815 | tartrate transporter | 2.372939979 | 7.40985E-11 |
| FGRAMPH1_01G23155 | hypothetical protein | 2.369185552 | 3.91097E-08 |
| FGRAMPH1_01G23285 | hypothetical protein | 2.369089952 | 4.12392E-32 |
| FGRAMPH1_01G26395 | aspartate aminotransferase | 2.365206918 | 3.95132E-24 |
| FGRAMPH1_01G17957 | hypothetical protein | 2.364460923 | 5.18416E-23 |
| FGRAMPH1_01G21519 | Cysteine dioxygenase | 2.364238204 | 2.4723E-09 |

| | | | |
|---|---|---|---|
| FGRAMPH1_01G04919 | hypothetical protein | 2.363990093 | 2.99057E-29 |
| FGRAMPH1_01G01349 | hypothetical protein | 2.363070834 | 1.36838E-27 |
| FGRAMPH1_01G22017 | hypothetical protein | 2.357898252 | 0.000238317 |
| FGRAMPH1_01G13411 | hypothetical protein | 2.357700808 | 2.03163E-19 |
| FGRAMPH1_01G06707 | sterol O-acyltransferase | 2.355431279 | 3.58162E-10 |
| FGRAMPH1_01G17757 | betaine aldehyde dehydrogenase | 2.355359742 | 2.43861E-25 |
| FGRAMPH1_01G01275 | hypothetical protein | 2.353430751 | 3.4624E-36 |
| FGRAMPH1_01G17531 | Sphingoid long-chain base transporter RSB1 | 2.345546676 | 8.64362E-26 |
| FGRAMPH1_01G03923 | hypothetical protein | 2.341709138 | 1.07922E-12 |
| FGRAMPH1_01G08279 | transmembrane 53-a | 2.339070244 | 7.58503E-10 |
| FGRAMPH1_01G08371 | hypothetical protein | 2.337971291 | 8.12386E-12 |
| FGRAMPH1_01G15349 | hypothetical protein | 2.333199409 | 4.83437E-20 |
| FGRAMPH1_01G05231 | hypothetical protein FPSE_00596 | 2.331894003 | 2.64192E-05 |
| FGRAMPH1_01G09859 | related to hexokinase | 2.331178246 | 9.12634E-13 |
| FGRAMPH1_01G17421 | hypothetical protein | 2.325268923 | 7.921E-11 |
| FGRAMPH1_01G00177 | integral membrane | 2.320327004 | 2.31634E-16 |
| FGRAMPH1_01G06477 | translation factor pelota | 2.317517835 | 3.80142E-10 |
| FGRAMPH1_01G10817 | hypothetical protein | 2.317500239 | 8.60872E-13 |
| FGRAMPH1_01G07967 | hypothetical protein | 2.316488671 | 7.29164E-13 |
| FGRAMPH1_01G14959 | hypothetical protein | 2.302675732 | 1.26578E-15 |
| FGRAMPH1_01G21623 | 2-amino-3-carboxymuconate-6-semialdehyde decarboxylase | 2.302268661 | 2.24893E-10 |
| FGRAMPH1_01G10573 | 3-methylcrotonyl- carboxylase alpha subunit | 2.301123259 | 4.78332E-41 |
| FGRAMPH1_01G06449 | hypothetical protein | 2.297596431 | 2.79471E-08 |
| FGRAMPH1_01G11757 | mfs sp sugar:h+ symporter | 2.286030261 | 6.37339E-19 |
| FGRAMPH1_01G25249 | 3-hydroxybenzoate 6-hydroxylase | 2.285534344 | 1.75248E-06 |
| FGRAMPH1_01G28013 | hypothetical protein | 2.283171234 | 1.17757E-05 |
| FGRAMPH1_01G19975 | tyrosine phosphatase 2 | 2.282826148 | 8.0933E-16 |
| FGRAMPH1_01G21449 | superoxide-generating nadph oxidase heavy chain subunit c | 2.277571395 | 6.88201E-10 |
| FGRAMPH1_01G02741 | hypothetical protein | 2.277340663 | 4.85224E-07 |
| FGRAMPH1_01G03539 | plasma membrane yro2 | 2.273089056 | 7.31075E-11 |
| FGRAMPH1_01G07277 | geranylgeranyl pyrophosphate synthetase | 2.272231722 | 5.45269E-08 |
| FGRAMPH1_01G14131 | aat family amino acid transporter | 2.269441283 | 2.02426E-11 |
| FGRAMPH1_01G06575 | arrestin containing | 2.267568748 | 3.25504E-14 |
| FGRAMPH1_01G24933 | hypothetical protein | 2.265129988 | 1.97451E-33 |
| FGRAMPH1_01G06959 | hypothetical protein | 2.264338769 | 1.36946E-06 |
| FGRAMPH1_01G08357 | endoribonuclease L-PSP | 2.262132952 | 3.81436E-08 |
| FGRAMPH1_01G08177 | Early growth response 1-B | 2.255173695 | 1.17449E-05 |
| FGRAMPH1_01G13061 | alcohol dehydrogenase Bli-4 | 2.250201674 | 8.03455E-10 |
| FGRAMPH1_01G03479 | related to rna-binding fus tls | 2.247795974 | 4.06975E-37 |
| FGRAMPH1_01G10409 | 2-oxoisovalerate dehydrogenase subunit beta | 2.246850138 | 1.12689E-15 |
| FGRAMPH1_01G21713 | hypothetical protein | 2.245277985 | 5.36155E-11 |

| | | | |
|---|---|---|---|
| FGRAMPH1_01G18257 | hypothetical protein | 2.243135626 | 1.61685E-15 |
| FGRAMPH1_01G01945 | ctr copper transporter family | 2.242890031 | 4.81597E-27 |
| FGRAMPH1_01G13919 | hypothetical protein | 2.228706261 | 1.73327E-06 |
| FGRAMPH1_01G15813 | triacylglycerol lipase | 2.227740567 | 1.37217E-05 |
| FGRAMPH1_01G23339 | hypothetical protein | 2.227722184 | 8.10654E-16 |
| FGRAMPH1_01G15061 | fructose-2,6-bisphosphatase | 2.226129787 | 1.68974E-08 |
| FGRAMPH1_01G00001 | hypothetical protein | 2.222289817 | 7.18326E-05 |
| FGRAMPH1_01G12697 | long-chain acyl- synthetase | 2.221990602 | 1.11956E-06 |
| FGRAMPH1_01G06447 | hypothetical protein | 2.217217222 | 3.67886E-09 |
| FGRAMPH1_01G17365 | hypothetical protein | 2.216894607 | 2.37012E-17 |
| FGRAMPH1_01G05343 | catalase | 2.212487107 | 9.08371E-05 |
| FGRAMPH1_01G12119 | o-acetyltransferase cas1 | 2.210682192 | 4.44012E-08 |
| FGRAMPH1_01G17743 | major facilitator superfamily transporter | 2.203340003 | 3.08308E-37 |
| FGRAMPH1_01G13617 | transmembrane amino acid transporter | 2.200976091 | 3.62558E-09 |
| FGRAMPH1_01G17367 | hypothetical protein | 2.198364563 | 1.91134E-11 |
| FGRAMPH1_01G26651 | hypothetical protein | 2.196628191 | 1.911E-07 |
| FGRAMPH1_01G06475 | hypothetical protein | 2.193619898 | 6.63179E-08 |
| FGRAMPH1_01G07905 | hypothetical protein | 2.188951335 | 3.48743E-05 |
| FGRAMPH1_01G05871 | hypothetical protein | 2.187448681 | 1.50199E-10 |
| FGRAMPH1_01G13665 | hypothetical protein | 2.186017663 | 9.51103E-09 |
| FGRAMPH1_01G14483 | hypothetical protein | 2.18546907 | 9.7696E-19 |
| FGRAMPH1_01G20215 | rna 3 -terminal phosphate cyclase | 2.183741573 | 1.52751E-14 |
| FGRAMPH1_01G12863 | auxin efflux carrier superfamily | 2.181206901 | 5.93398E-09 |
| FGRAMPH1_01G20985 | TOB3 (member of AAA-ATPase family) | 2.18072113 | 0.000153556 |
| FGRAMPH1_01G04363 | unnamed protein product | 2.179463516 | 2.15516E-06 |
| FGRAMPH1_01G03639 | hypothetical protein | 2.177655105 | 8.016E-06 |
| FGRAMPH1_01G02211 | hypothetical protein | 2.174823579 | 4.77078E-06 |
| FGRAMPH1_01G27899 | transcriptional regulatory | 2.174083613 | 2.1128E-10 |
| FGRAMPH1_01G21055 | lysine n-acyltransferase | 2.17031314 | 1.40599E-05 |
| FGRAMPH1_01G06295 | Beta,beta-carotene 15,15 -monooxygenase | 2.169336635 | 1.07782E-14 |
| FGRAMPH1_01G28053 | AAA family | 2.168777462 | 8.34201E-22 |
| FGRAMPH1_01G23197 | alpha beta fold family | 2.16810927 | 4.5413E-15 |
| FGRAMPH1_01G12571 | integral membrane PTH11 | 2.166165888 | 5.03541E-17 |
| FGRAMPH1_01G05759 | ca2+-transporting atpase | 2.158607144 | 2.31218E-18 |
| FGRAMPH1_01G11897 | msf membrane transporter | 2.158408681 | 2.4693E-21 |
| FGRAMPH1_01G10481 | amidophosphoribosyltransferase | 2.1569025 | 5.06123E-05 |
| FGRAMPH1_01G08355 | D-amino-acid oxidase | 2.156873908 | 1.38334E-08 |
| FGRAMPH1_01G02047 | v-type proton atpase subunit a | 2.156125442 | 1.54237E-10 |
| FGRAMPH1_01G10039 | pho85 cyclin-1 | 2.15571815 | 2.35864E-08 |
| FGRAMPH1_01G13065 | hypothetical protein | 2.154343881 | 1.44054E-07 |
| FGRAMPH1_01G01685 | hypothetical protein | 2.153796395 | 3.46777E-07 |
| FGRAMPH1_01G22469 | aat family amino acid transporter | 2.150918067 | 4.64726E-14 |

| | | | |
|---|---|---|---|
| FGRAMPH1_01G27703 | hypothetical protein | 2.148901288 | 1.96323E-08 |
| FGRAMPH1_01G10139 | isovaleryl-CoA dehydrogenase | 2.14462629 | 1.60869E-24 |
| FGRAMPH1_01G03789 | hypothetical protein | 2.140012988 | 1.23763E-21 |
| FGRAMPH1_01G26009 | hypothetical protein | 2.139527184 | 5.24335E-26 |
| FGRAMPH1_01G02281 | hypothetical protein | 2.139315062 | 9.89374E-23 |
| FGRAMPH1_01G19809 | hypothetical protein | 2.138043807 | 2.00792E-07 |
| FGRAMPH1_01G17543 | hypothetical protein | 2.136254991 | 6.8043E-18 |
| FGRAMPH1_01G27797 | mitochondrial phosphate carrier | 2.134781705 | 1.41149E-15 |
| FGRAMPH1_01G04771 | arylacetamide deacetylase | 2.132045425 | 2.28313E-33 |
| FGRAMPH1_01G06343 | unnamed protein product | 2.131724485 | 3.03178E-32 |
| FGRAMPH1_01G25119 | short-chain dehydrogenase | 2.126735708 | 0.000583992 |
| FGRAMPH1_01G27465 | hypothetical protein | 2.119203927 | 1.06225E-06 |
| FGRAMPH1_01G15391 | hypothetical protein | 2.118341058 | 1.03127E-13 |
| FGRAMPH1_01G15615 | hypothetical protein | 2.115689702 | 8.26655E-14 |
| FGRAMPH1_01G02017 | hypothetical protein | 2.113441745 | 8.40466E-22 |
| FGRAMPH1_01G17369 | hypothetical protein | 2.112238897 | 0.000236767 |
| FGRAMPH1_01G05803 | hypothetical protein | 2.107872057 | 2.25752E-13 |
| FGRAMPH1_01G01241 | hypothetical protein | 2.106310395 | 3.43319E-24 |
| FGRAMPH1_01G12421 | glycerol-3-phosphate dehydrogenase | 2.09829225 | 2.14963E-15 |
| FGRAMPH1_01G10137 | methylcrotonoyl-CoA carboxylase subunit beta | 2.097069796 | 1.43284E-11 |
| FGRAMPH1_01G02555 | 2-oxoisovalerate dehydrogenase E2 component (dihydrolipoyl transacylase) | 2.094586767 | 4.41606E-13 |
| FGRAMPH1_01G06565 | hypothetical protein | 2.088318168 | 1.46616E-09 |
| FGRAMPH1_01G20169 | unnamed protein product | 2.08362447 | 3.53558E-24 |
| FGRAMPH1_01G26493 | polyamine transporter 2 | 2.081544973 | 2.19422E-17 |
| FGRAMPH1_01G08295 | LITAF-like zinc finger domain-containing | 2.080123616 | 1.4152E-17 |
| FGRAMPH1_01G02149 | hypothetical protein | 2.076637746 | 5.99645E-05 |
| FGRAMPH1_01G17749 | l-ornithine n5-oxygenase | 2.072532497 | 6.91352E-21 |
| FGRAMPH1_01G06589 | hypothetical protein | 2.07185632 | 2.60591E-17 |
| FGRAMPH1_01G08361 | hypothetical protein | 2.060698996 | 4.76674E-07 |
| FGRAMPH1_01G10455 | hypothetical protein | 2.05839622 | 1.1479E-14 |
| FGRAMPH1_01G10127 | Regulator of rDNA transcription 8 | 2.056087555 | 2.03956E-14 |
| FGRAMPH1_01G07239 | hypothetical protein | 2.054804857 | 3.38936E-05 |
| FGRAMPH1_01G08631 | hypothetical protein | 2.052753708 | 1.63509E-07 |
| FGRAMPH1_01G05529 | hypothetical protein | 2.049883712 | 3.80104E-12 |
| FGRAMPH1_01G04455 | hypothetical protein | 2.049634921 | 0.000628279 |
| FGRAMPH1_01G01261 | hypothetical protein | 2.048509223 | 3.55295E-06 |
| FGRAMPH1_01G13395 | salicylate hydroxylase | 2.046330728 | 4.58467E-06 |
| FGRAMPH1_01G04101 | hypothetical protein | 2.045827639 | 2.76615E-07 |
| FGRAMPH1_01G01129 | hypothetical protein | 2.042834792 | 7.71516E-05 |
| FGRAMPH1_01G22647 | hypothetical protein | 2.042401326 | 0.000844794 |
| FGRAMPH1_01G02853 | ring finger b | 2.040484685 | 1.24978E-33 |

| | | | |
|---|---|---|---|
| FGRAMPH1_01G04241 | acetyl-coenzyme A synthetase | 2.039146276 | 1.82506E-06 |
| FGRAMPH1_01G06839 | transcriptional activator protein acu-15 | 2.039080606 | 1.02941E-08 |
| FGRAMPH1_01G12837 | chitin synthase 1 | 2.038109303 | 8.83005E-09 |
| FGRAMPH1_01G08381 | unnamed protein product | 2.036346809 | 1.28646E-12 |
| FGRAMPH1_01G22635 | acetyl-CoA carboxylase | 2.036274597 | 1.37782E-13 |
| FGRAMPH1_01G26081 | hypothetical protein | 2.035754503 | 3.47882E-16 |
| FGRAMPH1_01G13133 | NADH-cytochrome b5 reductase 1 | 2.031908106 | 0.000423425 |
| FGRAMPH1_01G18255 | unnamed protein product | 2.031176135 | 2.78474E-14 |
| FGRAMPH1_01G03517 | related to D-ribulokinase | 2.030665607 | 8.00026E-08 |
| FGRAMPH1_01G14957 | hypothetical protein | 2.029602193 | 2.48058E-06 |
| FGRAMPH1_01G25333 | quinate permease | 2.02195559 | 2.25285E-16 |
| FGRAMPH1_01G01761 | hypothetical protein | 2.021686729 | 1.58331E-14 |
| FGRAMPH1_01G27919 | hypothetical protein | 2.020996623 | 0.000870582 |
| FGRAMPH1_01G10711 | hypothetical protein | 2.017744357 | 1.30855E-05 |
| FGRAMPH1_01G13423 | Catechol 1,2-dioxygenase | 2.013943032 | 6.18174E-18 |
| FGRAMPH1_01G17151 | tetracycline resistance TCR1 | 2.007502162 | 2.88238E-07 |
| FGRAMPH1_01G07577 | plasma membrane proteolipid 3 | 2.004193642 | 9.95027E-22 |
| FGRAMPH1_01G20719 | related to DUF895 domain membrane | 2.004013145 | 1.13396E-09 |
| FGRAMPH1_01G01127 | hypothetical protein | 2.002744035 | 1.77623E-09 |
| FGRAMPH1_01G03505 | plasma membrane ATPase | 2.001398427 | 3.22929E-18 |
| FGRAMPH1_01G06971 | hypothetical protein | 2.000738096 | 0.000272557 |

**Supplementary Data 2. Identification of *F. graminearum* homologs of conserved fungal GlcNAc catabolic genes and their expression in the *ZEB2OE* strain.**

| Predicted function | Query gene | Identities | Gene ID | Annotation | *ZEB2OE*/WT log2(FC) | P-adj |
|---|---|---|---|---|---|---|
| GlcNAc transporter | Ngt | 69% | FGRAMPH1_01G20719 | related to DUF895 domain membrane | 2.004013145 | 1.13396E-09 |
| | | 59% | FGRAMPH1_01G09423 | DUF895 domain membrane | 1.435598624 | 0.136618636 |
| | | 58% | FGRAMPH1_01G25977 | probable DUF895 domain membrane | 1.676201339 | 1.3705E-20 |
| | | 29% | FGRAMPH1_01G25651 | DUF895 domain membrane | -1.083648381 | 0.00015758 |
| | | 27% | FGRAMPH1_01G05179 | DUF895 domain membrane | -0.438774612 | 0.516972924 |
| GlcNAc-hexokinase | Hxk | 47% | FGRAMPH1_01G09859 | related to hexokinase | 2.331178246 | 9.12634E-13 |
| | | 28% | FGRAMPH1_01G26225 | related to hexokinase | -1.810757171 | 3.39963E-17 |
| | | 27% | FGRAMPH1_01G10791 | hexokinase | -0.40092224 | 0.39239941 |
| | | 26% | FGRAMPH1_01G01281 | hexokinase | -0.684262978 | 0.007274907 |
| | | 24% | FGRAMPH1_01G11851 | hexokinase | -0.134626678 | 0.793227957 |
| | | 25% | FGRAMPH1_01G25401 | probable glucokinase | -0.854306581 | 0.0731267 |
| GlcNAc-6-phosphate deacetylse | Nag | 57% | FGRAMPH1_01G09851 | n-acetylglucosamine-6-phosphate deacetylase | 4.734823396 | 1.22031E-60 |
| | | 23% | FGRAMPH1_01G12759 | beta-glucosidase M | -1.352012959 | 1.78989E-05 |
| | | 22% | FGRAMPH1_01G16717 | beta-glucosidase g | -0.210327692 | 0.618780413 |
| | | 26% | FGRAMPH1_01G00457 | beta-glucosidase | 0.241582179 | 0.369459159 |
| Glucosamine-6-phosphate deaminase | Dam | 47% | FGRAMPH1_01G09857 | glucosamine-6-phosphate isomerase | 3.775410358 | 1.1251E-115 |
| | | 50% | FGRAMPH1_01G25397 | glucosamine-6-phosphate isomerase | -0.781366228 | 0.085435385 |
| Ndt80-like transcription factor | Ron | 42% | FGRAMPH1_01G09855 | acid phosphatase | -0.127198161 | 0.466412504 |
| | | 34% | FGRAMPH1_01G24101 | acid phosphatase | 0.466475181 | 0.359174296 |

**Supplementary Data 3. Strains used in this study.**

| Strains | Description | Reference |
|---|---|---|
| Z-3639 | *Fusarium graminearum* wild-type strain | [1] |
| *ZEB2OE* | *ZEB2*-overexpression strain | [2] |
| Δ*zeb2* | *ZEB2* knockout mutant | [2] |
| Δ*ron* | *RON* knockout mutant | This study |
| Δ*ngt* | *NGT* knockout mutant | This study |
| Δ*hxk* | *HXK* knockout mutant | This study |
| Δ*nag* | *NAG* knockout mutant | This study |
| Δ*dam* | *DAM* knockout mutant | This study |
| *NAGcom* | *NAG*-complemented strain | This study |
| *NAGOE* | *NAG*-overexpressed strain | This study |
| Δ*chs7* | *CHS7* knockout mutant | [3] |
| $P_{FCR1}$-*FgNAG* | *FgNAG* conditional-expression strain | This study |
| $P_{FCR1}$-*FgNAG*; *ZEB2LOE* | *ZEB2L*-overexpressed strain in the $P_{FCR1}$-*FgNAG* background | This study |
| *ZEB2OE*; *NAGOE* | *NAG*-overexpressed strain in the *ZEB2OE* background | This study |

**Supplementary Data 4. Primers used in this study.**

| Primer name | Sequence (5'–3') | Purpose |
|---|---|---|
| FgRON_5F | GTATGGAGGTCAGGACGAGGTT | |
| FgRON_5N | ACGGAAAGTAAGCTGAAGTGAGTG | |
| FgRON_5R | GCACAGGTACACTTGTTTAGAGGGAACCGTCAACGCCTCAT | |
| FgRON_3F | CCTTCAATATCATCTTCTGTCGTGAATTATGAATTGTTGGAAAAGCA | *RON* gene deletion |
| FgRON_3N | AGGTCAAGACAGTCAAGGCAAG | |
| FgRON_3R | AAGTGCGTCGAACAGGGC | |
| FgRON with 5F | GAATTGTCGAGTCGAATATGAGAAG | |
| FgNGT_5F | AGCCAATGCTGCTGTGTTTCA | |
| FgNGT_5N | ACCGCCACTAAAATTTGATGCT | |
| FgNGT_5R | GCACAGGTACACTTGTTTAGAGTGTGTGTGTGTGTGTGTGTGTA | |
| FgNGT_3F | CCTTCAATATCATCTTCTGTCGCGTTTTGAAGTGTACACATTGGA | *NGT* gene deletion |
| FgNGT_3N | GTCATCACTATTTATCGTCACCATC | |
| FgNGT_3R | GCGAGACTGCCAGCCGAC | |
| FgNGT with 5F | GTAAAGAGCTGTGTTGGCGTGAT | |
| FgHXK_5F | CGCACTTTCGGCGATACCAT | |
| FgHXK_5N | GAAGATGCAATAGATGGGATGACC | |
| FgHXK_5R | GCACAGGTACACTTGTTTAGAGAAAGGAGTGGGGGTTATGAAAG | |
| FgHXK_3F | CCTTCAATATCATCTTCTGTCGTTTTGTCTGAACATTTGGAGTAGAA | *HXK* gene deletion |
| FgHXK_3N | GTTTGGCTGGTTAGTTGTGTGTAG | |
| FgHXK_3R | TAAGAATCTCAGGGTCGGGCA | |
| FgHXK with 5F | TAGGCAGAGGCCCGTCAGAT | |
| FgNAG_5F | TTGACATGAATCCAATTGTCTCTAA | |
| FgNAG_5N | CCGGTCGAGATCGTGCCAT | |
| FgNAG_5R | GCACAGGTACACTTGTTTAGAGATTGAAGGATTGTAGGAGTGGTCT | |
| FgNAG_3F | CCTTCAATATCATCTTCTGTCGAGTGATGCTGGTTGGTGCTC | *NAG* gene deletion |
| FgNAG_3N | CAGTAGGCAGCCCCGACAT | |
| FgNAG_3R | AAACGTACAGCAGCGACTCATC | |
| FgNAG with 5F | GCTGACTTGTGATGGTTGGTATG | |
| FgDAM_5F | AACGTTGCGTGTGAGTGCCC | |
| FgDAM_5N | CCATAAAAATGTTAGTCCCAAGTG | |
| FgDAM_5R | GCACAGGTACACTTGTTTAGAGAGTGTTGTTGAATTGCAAGGTGG | *DAM* gene deletion |
| FgDAM_3F | CCTTCAATATCATCTTCTGTCGAGGAATATGAATCGAGGAGGAAG | |

| | | |
|---|---|---|
| FgDAM_3N | CGATATCCTCCTGCCAATCTTC | |
| FgDAM_3R | GACCCCTCTATCTCAACCACTGC | |
| FgDAM with 5F | GAGTTGGAGAGGATGGTGTCGTAG | |
| Gen with 5F | GTTGCCTAGTGAATGCTCCGTAACA | Geneticin resistance cassette screening |
| Ron qRT-F | TCTTCGTAGCCGAGGATGTATTC | qRT-PCR analysis |
| Ron qRT-R | ATGTGGGTTGGCATTCTTCC | |
| Ngt qRT-F | CTCTGGTACGCCTCGCCTAAA | |
| Ngt qRT-R | GTAAAGAGCTGTGTTGGCGTGAT | |
| Hxk qRT-F | TTCGACCCTCAGAGTAGCATTG | |
| Hxk qRT-R | TAGGCAGAGGCCCGTCAGA | |
| Nag qRT-F | CCAACTGTCGATTGCTGAAAG | |
| Nag qRT-R | GCCATGTTATCGAGTAGGGTTGA | |
| Dam qRT-F | CGACCCCAACTCATATCATTCATT | |
| Dam qRT-R | GCGGCTGGCAAGACTGG | |
| HYG-F1 | GGCTTGGCTGGAGCTAGTGGAGG | *HYG-pFCR1* PCR construct |
| pFCR1-R1 | GGTGGCAGTTTCTGACATGCC | |
| HYF-pFCR1-5N | GCGTCGGTTTCCACTATCG | |
| HYG-pFCR1-3N | AGCGGTCATTGACTGGAGC | |
| NAG pFCR1-5F | CCGGTCGAGATCGTGCCAT | *NAG* promoter replacement |
| NAG pFCR1-5R | CCTCCACTAGCTCCAGCCAAGCCAAAACTGAGGGCGA ATGGTATTATAC | |
| NAG pFCR1-3F | GGCATGTCAGAAACTGCCACCATGCCCATCGCAGTCT CG | |
| NAG pFCR1-3R | ATAAGCCCCATCAGGAAGACC | |
| NAG hyg_5N | TATTTCGGTCTTTTCTGATGGTG | |
| NAG-pFCR1-3N | CGATTTTGATGGTTGTGGGG | |
| NAG com-F | TATAGGGCGAATTGGGTACTCAAATTGGTTTTGACATG AATCCAATTGTCTCTAA | *NAG* complementation & overexpression |
| NAGOE-F | TTTCGTAGGAACCCAATCTTCAAAATGCCCATCGCAGT CTCGCCAAC | |
| NAG-R | CCCGGTGAACAGCTCCTCGCCCTTGCTCACCTTCAGCT TACTTTCCGTGGCA | |
| Rp27 with F | GGGAACTTTTTCTCACAGACACAA | Clone seqeuncing |
| Native seqF-1 | CCAGCTGGCGAAAGGGG | |
| GFP-R | GTGGTGCAGATGAACTTCA | |
| ZEB2LOE_F | TTTCGTAGGAACCCAATCTTCAAAATGCAATCTACCGA GTCGTTCCAT | *ZEB2L* overexpression construct with a mutated *ZEB2S* start codon |
| ZEB2L_mutation _with_F | CATGGTGCCCGAGGAGAAAcTGAGTCGTATTCTCAAG GT | |
| ZEB2L_mutation _with_R | ACCTTGAGAATACGACTCAgTTTCTCCTCGGGCACCAT G | |
| ZEB2LOE_R | CCCGGTGAACAGCTCCTCGCCCTTGCTCACTTAGCATT GAGCTGGATTGAAG | |

### References

1 Bowden, R. L. & Leslie, J. F. Sexual recombination in *Gibberella zeae*. *Phytopathology* **89**, 182–188 (1999).

2 Park, A. R. *et al.* Autoregulation of *ZEB2* expression for zearalenone production in *Fusarium graminearum*. *Mol. Microbiol.* **97**, 942–956 (2015).

3 Kim, J.-E. *et al. Gibberella zeae* chitin synthase genes, GzCHS5 and GzCHS7, are required for hyphal growth, perithecia formation, and pathogenicity. *Curr. Genet.* **55**, 449–459 (2009).
